# Deficits in neuronal architecture but not over-inhibition are main determinants of reduced neuronal network activity in a mouse model of overexpression of *Dyrk1A*

**DOI:** 10.1101/2023.03.09.531874

**Authors:** Linus Manubens-Gil, Meritxell Pons-Espinal, Thomas Gener, Inmaculada Ballesteros-Yañez, María Martínez de Lagrán, Mara Dierssen

## Abstract

Abnormal dendritic arbors, dendritic spine “dysgenesis” and excitation inhibition imbalance are main traits assumed to underlie impaired cognition and behavioral adaptation in intellectual disability. However, how these modifications actually contribute to functional properties of neuronal networks, such as signal integration or storage capacity is unknown. Here, we used a mouse model overexpressing *Dyrk1A* (Dual-specificity tyrosine [Y]-regulated kinase), one of the most relevant Down syndrome (DS) candidate genes, to gather quantitative data regarding hippocampal neuronal deficits produced by the overexpression of *Dyrk1A* in mice (TgDyrk1A; TG). TG mice showed impaired hippocampal recognition memory, altered excitation-inhibition balance and deficits in hippocampal CA1 LTP. We also detected for the first time that deficits in dendritic arborization in TG CA1 pyramidal neurons are layer-specific, with a reduction in the width of the *stratum radiatum*, the postsynaptic target site of CA3 excitatory neurons, but not in the *stratum lacunosum-moleculare*, which receives temporo-ammonic projections. To interrogate about the functional impact of layer-specific TG dendritic deficits we developed tailored computational multicompartmental models. Computational modelling revealed that neuronal microarchitecture alterations in TG mice lead to deficits in storage capacity, altered the integration of inputs from entorhinal cortex and hippocampal CA3 region onto CA1 pyramidal cells, important for coding place and temporal context and on connectivity and activity dynamics, with impaired the ability to reach high γ oscillations. Contrary to what is assumed in the field, the reduced network activity in TG is mainly contributed by the deficits in neuronal architecture and to a lesser extent by over-inhibition. Finally, given that therapies aimed at improving cognition have also been tested for their capability to recover dendritic spine deficits and excitation-inhibition imbalance, we also tested the short- and long-term changes produced by exposure to environmental enrichment (EE). Exposure to EE normalized the excitation inhibition imbalance and LTP, and had beneficial effects on short-term recognition memory. Importantly, it produced massive but transient dendritic remodeling of hippocampal CA1, that led to recovery of high γ oscillations, the main readout of synchronization of CA1 neurons, in our simulations. However, those effects where not stable and were lost after EE discontinuation. We conclude that layer-specific neuromorphological disturbances produced by Dyrk1A overexpression impair coding place and temporal context. Our results also suggest that treatments targeting structural plasticity, such as EE, even though hold promise towards improved treatment of intellectual disabilities, only produce temporary recovery, due to transient dendritic remodeling.

## INTRODUCTION

Down syndrome (DS; trisomy 21) is the most frequent genetic cause of intellectual disability. DS learning and memory deficits are assumed to be caused by synaptic dysfunction, and impaired activity-dependent plasticity^1^. Among several human chromosome 21 (I21) genes, overexpression of *Dyrk1A*^2^ (Dual-specificity tyrosine [Y]-regulated kinase) in transgenic mice (TgDyrk1A; TG) impairs hippocampal-dependent learning and memory^3,4^, and is sufficient to recapitulate DS neuronal architecture deficits and reduced γ oscillations in the cerebral cortex^5^. Although it has long been assumed that those microstructural alterations along with the over-inhibition^6,7^ of the network account for the specific cognitive deficits in DS, how those modify network activity is unknown. Although function cannot be directly estimated from structure, it can be inferred by statistical, and biophysical models that help to translate brain structure to brain function. For this reason, we developed tailored computational multicompartmental models to assess the impact of microstructural deficits on connectivity and activity dynamics, and specifically, on the integration of inputs from entorhinal cortex and hippocampal CA3 region onto CA1 pyramidal cells, important for coding place and temporal context

Moreover, therapies aimed at improving cognition have also been tested for their capability to recover structural deficits. The cognitive benefits of environmental enrichment (EE) are well substantiated. In the Ts65Dn partial trisomic DS mouse model^8^, one month of exposure to EE was sufficient to reduce GABAergic release^9^, and to rescue hippocampal-dependent cognitive and neuronal impairments^10^. The most prominent effects of EE in trisomic mice were dendritic branching and spine density rearrangements^11^ mainly in the hippocampus. However, it is unknown how those structural changes translate into changes in functional properties. It is known that EE increases the theta-associated γ oscillation power in the hippocampal CA1 in rats^12^, which is expressed most prominently in the *stratum radiatum*, where axons from bilateral CA3 pyramidal cells make synaptic connections. This is interesting because EE normalizes DYRK1A kinase activity in the hippocampus^8^, and TG mice overexpressing *Dyrk1A* show decreased firing rate and gamma frequency power in the prefrontal network. Moreover, reducing the amplitude and coordinating γ oscillations with optogenetics ameliorates behavioral deficits in an Alzheimer’s disease mouse model^13^. Thus, we here explored the specific contribution of DYRK1A to the EE improvements.

In summary, we here tested the hypothesis that synaptic and structural neuronal defects lead to cognitive defects in TG mice by impairing hippocampal CA1 integration of CA3 and entorhinal cortex (EC) inputs. We identified clear structural deficits in TgDyrk1A neurons along with altered excitation inhibition balance, that were temporarily rescued by EE through massive dendritic remodeling specifically affecting the distal dendritic segment of pyramidal CA1 neurons. To understand the functional implications of these alterations, we developed tailored co^15^mputational multicompartmental models of single neurons to assess the impact of layer-specific TG dendritic deficits on connectivity and activity dynamics.

## RESULTS

### Environmental enrichment (EE) improves cognitive performance, and partially rescues levels of DYRK1A kinase activity, dendritic complexity and spine morphology in young (2-months) TgDyrk1A

We tested the effects of genotype and EE using the novel object recognition paradigm in 2-month-old mice (2mo). TgDyrk1A (TG) non-enriched (NE) showed impaired hippocampal recognition memory, with reduced novel object discrimination (discrimination index; DI) compared to WT (Fig. 1a; two-way ANOVA genotype x treatment interaction F(1,117)=8.1, p=0.005; TGNE vs. WTNE p=0.05) with no genotype-dependent differences in total time of exploration in the training session (see Table 3 for a summary of group comparison in each figure; see Shiny app https://linusmg.shinyapps.io/EE_TgDyrk1A/ for interactive analysis). This impairment in DI was rescued by one month of EE (Fig. 1a; two-way ANOVA treatment effect F(1,117)=7.6, p=0.007; TGEE vs. TGNE p=0.002). EE did not show effects in WT mice (Fig 1a; WTEE vs. WTNE, N.S.). Hippocampal DYRK1A kinase activity showed a significant genotype-treatment interaction (Fig 1b; two-way ANOVA genotype-treatment interaction F(1,79)=5.8, p=0.02), but pairwise comparisons for WTNE vs. TGNE and TGEE vs. TGNE were not statistically significant.

**Figure 1.**
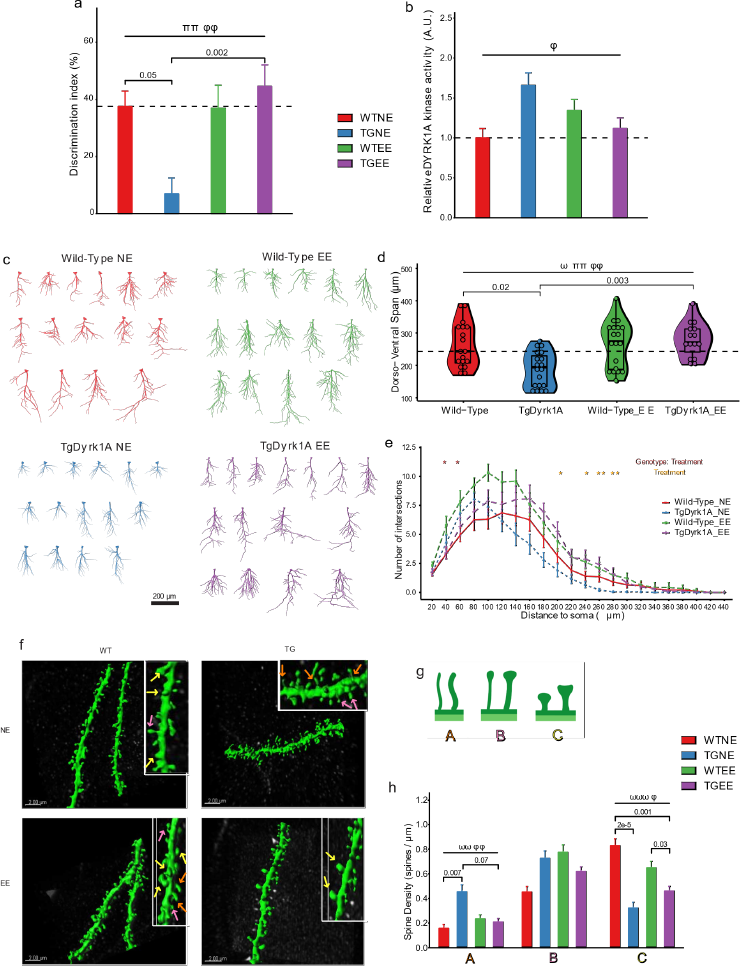
Environmental enrichment (EE) improves cognitive performance, and partially rescues levels of DYRK1A kinase activity, dendritic complexity and spine morphology in 2 month-old (2mo) TgDyrk1A (TG) **a** Discrimination index (%) during the test session of the NOR paradigm in wild type (WT) and TgDyrk1A (TG) mice reared for one month under non-enriched (NE) or environmentally enriched (EE) conditions. Number of animals analyzed: WTNE (2mo) = 14; TGNE (2mo) = 15; WTEE (2mo) = 16; TGEE (2mo) = 19. **b** Relative DYRK1A kinase activity in the hippocampus. Number of animals analyzed: WTNE (2mo) = 11; TGNE (2mo) = 9; WTEE (2mo) = 10; TGEE (2mo) = 10. For **a**-**b** Data are expressed as mean ± SEM. Two-way ANOVA treatment effect ππ p<0.01. Two-way ANOVA genotype-treatment interaction φφ p<0.01. **c** 2D projection of reconstructed apical dendritic trees in 2mo WT and TG, NE and EE. Scale bar = 200 μm. **d** Length of CA1 apical dendritic trees along the dorso-ventral axis (single cell measurement). Data are presented as an overlay of dot plots, boxplots and violin plots. Dotted line indicates the median value of the WTNE (2mo) group. Number of neurons/animals: WTNE (2mo) = 18/4; TGNE (2mo) = 19/4; WTEE (2mo) = 19/4; TGEE (2mo) = 15/3. Linear mixed effects: treatment effect ππ p<0.01. **e** Sholl analysis. Lines indicate mean ± SEM. Linear mixed effects, * p<0.05, ** p<0.01. **f** Representative confocal images showing CA1 apical dendritic spines in 2mo WT and TG mice under NE or EE conditions. Scale bar = 2 μm. Right upper insert corresponds to a higher magnification. Orange, pink and white arrows denote type A, type B and type C spines, respectively (see **g**). **g** Illustration of the morphological categories used to classify spines. **h** Density of each spine type. Data are expressed as mean ± SEM. Number of dendrite segments/neurons/animals analyzed: WTNE (2mo) = 19/19/4; TGNE (2mo) =16/12/4; WTEE (2mo) = 19/19/4; TGEE (2mo) = 16/9/4. Linear mixed effects: genotype effect ⍵⍵ p<0.01; ⍵⍵⍵ p<0.001; genotype x treatment interaction φ p<0.05; φφ p<0.01.

Reconstructions of pyramidal CA1 neurons in NE mice showed altered global neuronal architecture with a significant reduction in the dorso-ventral span (see Methods and Fig. S1), i.e. the length from cell body to apical tuft in TGNE mice (Figs. 1c and 1d; linear mixed effects genotype effect F(1,27.97)=4.61, p=0.04; TGNE vs. WTNE p=0.02). We also detected a reduction in the width of the *stratum radiatum* (*SR*), the postsynaptic target site of CA3 excitatory neurons, but not in the *stratum lacunosum-moleculare* (*SL*), which receives temporo-ammonic projections (Fig. S3a-c; linear mixed effects genotype effect for 2mo groups F(1,26.43)=15.26, p=6e-4; TGNE vs. WTNE p=0.003). EE rescued the reduction of dorso-ventral span of TG CA1 neurons (Fig. 1d; linear mixed effects treatment effect F(1,30.35)=10.80, p=0.003; genotype-treatment interaction F(1,30.43)=9.49, p=0.004; TGEE (2mo) vs. TGNE (2mo) p=0.003), but did not induce significant changes in WT neurons. We also found a significant dendritic remodeling of the dorso-ventral span upon EE (Fig. S3a-c; linear mixed effects genotype x treatment interaction for 2mo groups F(1,26.43)=5.67, p=0.03). However, post-hoc pairwise comparisons were not significant for either TGEE vs. TGNE nor WTEE vs. WTNE. The maximum path length, e.g. the length of the longest (primary) dendrite, was significantly reduced in TGNE neurons and recovered by EE treatment (Fig. S4a; linear mixed effects genotype x treatment interaction for 2mo groups F(1,28.19)=21.79, p=7e-5; treatment effect F(1,28.15)=13.49, p=0.001; TGNE vs. WTNE p=0.003; TGEE vs. TGNE p=6e-5), but again EE did not have significant effects on WT neurons.

Dendritic tree complexity was also reduced in TGNE CA1 neurons (Fig. 1c, 1e and S4b; Sholl area under the curve (AUC) linear mixed effects genotype effect for 2mo groups F(1,36.61)=5.99, p=0.02; TGNE vs. WTNE p=0.02). EE increased dendritic complexity in both TG and WT dendritic trees (Fig. 1e and Fig. S4b, linear mixed effects treatment effect for 2mo groups F(1,54.53)=12.07, p=0.001). The Sholl AUC was 52% larger in TGEE than in TGNE and 36% in WTEE compared to WTNE.

Finally, we did not observe differences in the total density of dendritic spines between genotypes (Fig. 1f and S3). However, TGNE mice showed reduced mature mushroom-like spines (Fig. 1f-h, linear mixed effects genotype effect F(1,24.69)=50.52, p=2e-7; TGNE vs. WTNE, Type C p=2e-5) along with increased thin immature spines (linear mixed effects genotype effect F(1,20.82)=8.29, p=0.009; TGNE vs. WTNE, Type A p=0.007). Regarding dendritic spines, the effects of EE were different in TG and WT. In TG neurons EE did not change total spine density but slightly increased mushroom-like spines (Fig. 1f, h; genotype-treatment interaction F(1,29.57)=5.23, p=0.03), at the expense of a significant reduction of thin spines (Fig. 1f, h; linear mixed effects genotype x treatment interaction F(1,22.88)=10.48, p=0.004; with a trend on TGEE vs. TGNE, type A p=0.07). In WT mice, EE did not produce any significant change nor any trend in the density of spines of any type.

In summary, EE rescued the cognitive impairment and hippocampal DYRK1A kinase activity in TG mice at 2mo of age. CA1 *SR* width defects were partially rescued by EE, which mainly increased dendritic complexity in both genotypes, whereas it possibly favored spine maturation and stabilization in TG mice.

### Deficits in LTP and excitation inhibition balance in young TG CA1 are rescued by EE

Extracellular field recordings on the Schaffer collateral-CA1 synapse of hippocampal slices measuring field excitatory postsynaptic potentials (fEPSPs) showed a significant genotype effect with reduced paired pulse facilitation (PPF, the ratio of a double-pulse stimulation fEPSP2/fEPSP1) in 2mo TG mice (Fig. 2a; two-way ANOVA genotype effect genotype effect F(1,105)=3.95, p=0.049), indicating an impairment of pre-synaptic function in TG mice. After TBS on the Schaffer collateral pathway, TGNE showed a significant deficit in hippocampal CA1 LTP (Fig. 2b, two-way ANOVA repeated measures, genotype effect F(1,43)=4.83, p<0.05). Upon EE, PPF was significantly increased in both genotypes (Fig. 2a; two-way ANOVA treatment effect F(1,105)=13.81, p=0.0003; TGEE vs. TGNE, p=0.001). Importantly, EE completely recovered the fEPSP slope (Fig. 2b; two-way ANOVA repeated measures treatment effect F(1,43)=3.76, p<0.05) and rescued the reduced LTP response in TG CA1.

**Figure 2.**
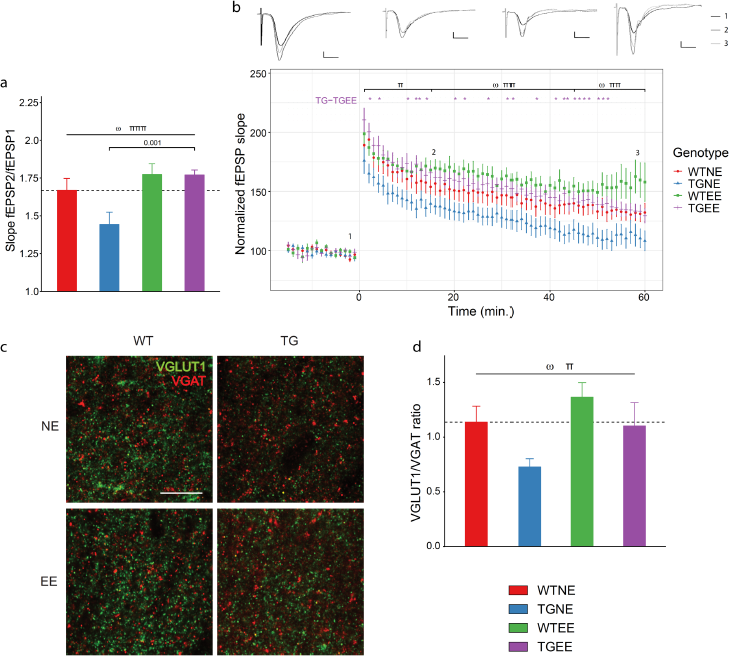
Deficits in LTP and excitation inhibition balance in TG CA1 are rescued by environmental enrichment. **a** Paired pulse facilitation (fEPSP2/fEPSP1) in the hippocampus of WT and TG mice reared under NE or EE conditions (slices/animals: WTNE (2mo) = 11/5; TGNE (2mo) = 11/5; WTEE (2mo) = 9/5; TGEE (2mo) = 16/5). Two-way repeated measures ANOVA genotype effect ⍵ p<0.05; treatment effect πππ p<0.001. **b** Long-term potentiation (LTP) in CA1 induced by theta burst stimulation (TBS) in CA3 under NE or EE conditions at 2 months of age (number of slice/animals: WTNE (2mo): 11/5; TGNE (2mo): 14/5; WTEE (2mo): 14/6; TGEE (2mo): 16/5). Upper panel: representation of the mean of 5 responses to a stimulation at time 1 (−5 min), 2 (15 min) and 3 (60 min). Vertical bar represents 500µV, horizontal bar represents 5ms. Lower panel: normalized fEPSP slope before and after TBS. Black horizontal bar represents 2 ms. Two-way repeated measures ANOVA genotype effect ⍵ p<0.05; treatment effect π p<0.05; ππ p<0.01. **c** Representative confocal images showing VGLUT1+ and VGAT+ puncta in CA1 *SR* hippocampal subfield. Scale bar = 10 mm. **d** Histogram showing the VGLUT1+ / VGAT+ ratio. Data are expressed as mean ± SEM. WTNE (2mo) n = 7; TGNE (2mo) n = 7; WTEE (2mo) n = 8; TGEE (2mo) n = 7. Two-way ANOVA genotype effect ⍵ p<0.05; treatment effect π p<0.05.

Given that impaired synaptic plasticity in DS may result from an excitation-inhibition imbalance^14,15^, we analyzed the ratio of excitatory (vesicular glutamate transporter 1, VGLUT1+) and inhibitory (vesicular GABA transporter, VGAT+) presynaptic puncta (Fig. 2c). TGNE 2mo mice showed a significantly reduced VGLUT1/VGAT ratio (Fig. 2d; two-way ANOVA genotype effect F(1,41)=5.81, p=0.02), indicating a bias towards inhibition in presynaptic transmission, due to reduced density of VGLUT1 puncta with no changes in VGAT density (Fig. 2c and 2d). EE increased VGLUT1/VGAT ratio in both genotypes, leading to a complete recovery of the excitatory-inhibitory imbalance in TGEE mice (Fig. 2c and 2d; two-way ANOVA treatment effect F(1,41)=4.67, p=0.04) due to the increase in VGLUT1 density (Fig. 2d).

### Connectivity repertoire and input-output frequency deficits in young TG mice are partially rescued by environmental enrichment

To infer whether the disturbances of dendritic architecture and spine density detected in 2mo TG CA1 pyramidal neurons may affect their topological properties, we quantified the repertoire of their possible connectivity patterns (Eq. 2, connectivity repertoire; Wen, 2009). To dissect the contribution of spine density on the connectivity repertoire, we calculated the number of contacts by assuming all experimental groups had the WT density of mature spines. When only accounting for the reduced dendritic span and complexity of TGNE neurons, the connectivity repertoire was not different from WTNE (Fig. S5a; linear mixed effects genotype effect for 2mo groups F(1,33.96)= 4.78, p=0.04; TGNE vs WTNE N.S.). However, when taking into account the reduction in mature spine density, it was significantly reduced in TGNE neurons (Fig. S6a and S5b; linear mixed effects genotype effect for 2mo groups F(1,35.88)= 20.69, p=6e-5; TGNE vs WTNE p=0.02). Upon EE, the increase in dendritic tree span and complexity (see Figs. 1d and 1e) detected in WTEE and TGEE neurons was sufficient to significantly increase the connectivity repertoire of TGEE neurons (Fig. S5a; linear mixed effects treatment effect F(1,46.01)= 13.29, p=0.0007; TGEE vs TGNE p=0.03). The increase in connectivity repertoire was also significant when we accounted for the increase of mature dendritic spines upon EE (Fig. S6a and S5b; linear mixed effects treatment effect for 2mo groups F(1,52.27)= 17.67, p=0.0001; TGEE vs TGNE p=0.049).

As reduced connectivity repertoire may affect neuronal signal integration^16^, we explored the input-output (IO) relation of individual CA1 pyramidal neurons using multi-compartmental models. Those models allowed simulating the complete tree spanning *SR* and *SL* (Figs. S6b, S7a and 3c) or the dendritic area covering *SR* alone (Figs S7b and 3d). There is a tendency of the CA1 region to phase lock with either CA3 or the medial entorhinal cortex at distinct frequencies, thus allowing to model input-output of *SR*, the postsynaptic target site of CA3 excitatory neurons, and *SL*, which receives temporo-ammonic projections, as both dendritic regions are differentially affected in the TGNE mice. We simulated low (40 Hz; Figs S8a and S8c) or high (100 Hz; Figs. S6d, S8b and S8c) frequency (LF and HF) inputs on each of the WT and TG virtual neuronal trees (n = 71), and modeled their output firing. Upon LF (40 Hz) inputs to the complete dendritic tree spanning *SR* and *SL*, TGNE neurons showed a significantly lower success rate of evoked spikes (82%) than WTNE (100%) (Fig. S8a; linear mixed effects genotype x treatment interaction for 2mo groups, F(1,39.84)= 7.47, p=0.009; TGNE vs WTNE p=0.04). As a consequence, the firing rate (averaged spikes/sec in all dendritic trees) of TGNE neurons was reduced (Fig. S6c; output firing rate TGNE (2mo) vs WTNE (2mo) p<0.05), but corresponded to low γ both in TGNE (32 Hz), and in WTNE (40 Hz).

When we simulated HF (100 Hz) inputs on the complete dendritic tree (*SR* and *SL*) in WTNE neurons, we detected the success rate of evoked spikes expected with HF ^17^ (51%; Fig. S8b), with an output firing rate at high γ (51 Hz). When simulating TGNE neurons with HF we detected a significantly lower success rate of evoked spikes (39%) than in WTNE (Fig. S8b and S7B, linear mixed effects genotype effect for 2mo groups F(1,29.86)= 18.98, p=1e-4; TGNE vs WTNE p=0.009). This led to a reduced output firing rate (39 Hz) in 2mo TG neurons, that did not reach high γ (Fig. S6c; IO frequency TGNE (2mo) vs WTNE (2mo) p<0.05 for LF input p<0.05 for HF input). In fact, when we modeled membrane potential (mV) in the multicompartmental simulations of representative dendritic trees covering *SR* and *SL,* TGNE neurons did not reach depolarization (Fig. S6b). See <u>Supplementary Text</u> and Figures S6-8 for results of simulating inputs on only *SR*, for which TGNE neurons responded in the β range.

EE had a strong impact on the success rate of evoked spikes and the output firing, so that both TGEE and WTEE neurons reach high γ output (>=50 Hz; Fig. S6c; IO frequency TGEE vs TGNE p<0.05 for LF input and p<0.05 for HF input; Fig. S6c and Fig S8a; success rate of evoked spikes upon LF stimulation of both *SR* and *SL* linear mixed effects treatment effect for 2mo groups F(1,39.49)= 7.55, p=0.009; TGEE vs. TGNE p=0.01; Fig. S8b; HF stimulation of both *SR* and *SL* linear mixed effects treatment effect for 2mo groups F(1,34.42)= 17.24 p=2e-4; TGEE vs. TGNE p=0.02). When only *SR* was stimulated with HF, TGEE neurons were not able to fire above β (see <u>Supplementary Text</u>)

### Deficits in network activity of young TG CA1 neurons are rescued by environmental enrichment

We next investigated whether the genotype- and EE-dependent changes in output firing could affect the synchronized activity of CA1 neurons. We simulated simultaneous HF inputs (100 Hz), to recapitulate the effect of a tetanic stimulation, on the whole population of reconstructed neurons (neurons/animals: WTNE (2mo) = 18/4; TGNE (2mo) = 19/4; WTEE (2mo) = 19/4; TGEE (2mo) = 15/3) and measured their averaged activity. For these simulations, we concentrated on the combined *SR* and *SL* input and we also introduced inhibitory feedback to explore the effect of the stimulation at the population level in a more realistic way (see Methods) by including in the model the over-inhibition detected in TGNE neurons and its recovery upon EE (Fig. 2c, d).

In all the simulations, we chose the proportion of inhibitory synapses based on the VGLUT/VGAT ratio measured in each experimental group. We then adjusted the proportion of active excitatory synapses of WTNE at 15% to recapitulate the physiological single neuron average firing rate upon 100 Hz stimulation of Schaffer collaterals, in the presence of inhibitory feedback (20 Hz^18^). To take into account variations in mature spine density in the experimental groups, we adjusted the density of excitatory synapses in TG and EE-treated neurons by multiplying the ratio of mature spines to WTNE.

We detected a decreased activity of TGNE neuronal population compared to WTNE (Fig. 3a, 3d and 3e; two-way ANOVA genotype effect F(1,32)=441.10, p=3e-20; TGNE vs. WTNE p=5e-11). As over-inhibition is considered a hallmark of DS pathology, to disentangle the impact of TGNE over-inhibition alone, we next simulated “healthy” morphology (simulating the morphology of WTNE neuron) but with TGNE inhibition (TGNE-Inh). This led to a 32% reduction, showing that the over-inhibition alone cannot completely explain the reduced TG network response (Fig. 3b, 3d and 3e; TGNE-Inh vs. TGNE (2mo) p=9e-3). Since we hypothesized that alterations in neuronal architecture should also contribute to a suboptimal network response, we simulated the “normal” inhibitory feedback (the proportion of VGAT puncta of WTNE neurons) in TGNE neurons to dissect the impact of neuron morphology alone. In these “normalized inhibition” conditions, at HF stimulation (100 Hz), TGNE neuron population already showed a 44% reduction in the power of the averaged activity (Figs. 3c, 3d and 3e; TGNE-Morph vs. TGNE (2mo) N.S.) indicating that the morphological changes are an important factor in the reduction of network activity.

**Figure 3.**
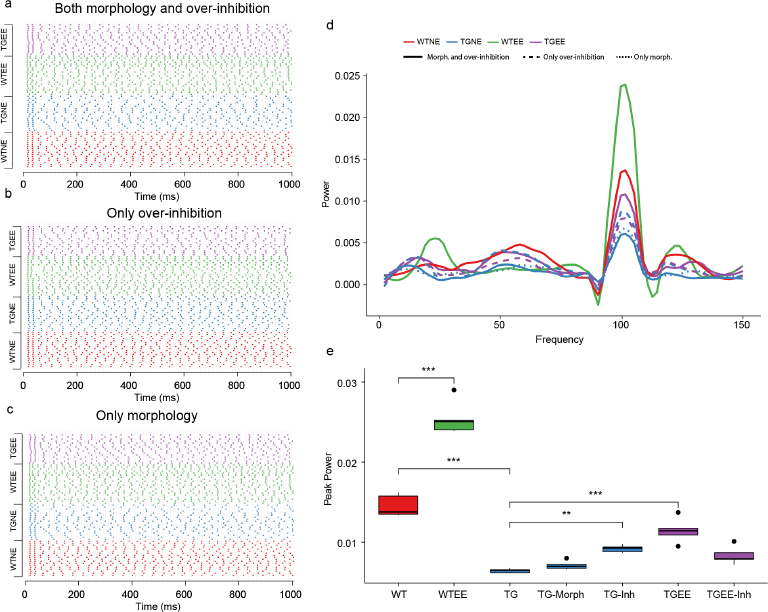
Deficits in network activity of TG CA1 neurons are rescued by environmental enrichment. Multi-compartmental models were generated for 2mo WT and TG neurons with a number of inhibitory synapses based on proportions known in WT CA1 neurons. **a-c** Spike raster plot of the simulated activity of CA1 neurons of 2mo WTNE (red), TGNE (blue) WTEE (green) and TGEE (purple). In a, b and c, each cell was stimulated with a 100 Hz input and with an inhibitory feedback loop. **a** Models were generated with TGNE and WTNE (2mo) morphologies for EE and NE groups and we modeled the number of inhibitory synapses based on measured VGLUT/VGAT ratio for 2mo TG and WT populations. Note the reduced firing in TG (2mo) neurons upon 100 Hz stimulation that is improved upon EE. **b** In this case, models were generated with only WTNE (2mo) morphologies for EE and NE groups and we modeled the number of inhibitory synapses based on measured VGLUT/VGAT ratio for 2mo TG and WT populations. **c** Models were generated for 2mo WT and TG neuron morphologies with a number of inhibitory synapses based on measured VGLUT/VGAT ratio. **d** Power spectra of spiking activity obtained averaged over 10 simulation repetitions for WTNE (2mo) and TGNE (2mo). Dashed lines show the results of simulations accounting only for over inhibition (TG-Inh are WT morphologies with TG inhibition; TGEE-Inh are TG morphologies with TGEE inhibition; corresponding to the raster plot in b) and solid lines represent simulations accounting for both morphological features and over inhibition (corresponding to the raster plot in a). **e** Peak power at 100 Hz for each of the neuron populations presented as boxplots. Two-way ANOVA, ** p<0.01, *** p<0.001. Number of neurons/animals: WTNE (2mo) = 18/4; TGNE (2mo) = 19/4; WTEE (2mo) = 19/4; TGEE (2mo) = 15/3.

We then explored the impact of EE on network activity, as it strongly improves both morphological and excitation-inhibition balance deficits in TG mice. At HF stimulation, the averaged activity of TGEE neurons was increased by 42% with respect to TGNE (Figs. 3a, 3d and 3e; treatment effect F(1,32)=230.04, p=4e-16; TGEE vs. TGNE p=5e-6). However, when simulating only the rescue of over-inhibition, TGEE only reached a 14% increase (Fig. 3b), showing again that morphology has a stronger impact on neuronal population response, as also confirmed by spectral analysis (Figs. 3d and 3e, dashed purple line is TG morphology with TGEE over-inhibition; TGEE-Inh vs. TGNE N.S.). Instead, confirming our hypothesis, the morphological changes in WT neurons upon EE treatment, led to 41% increased power (Figs. 3a, 3d and 3e; WTEE vs. WTNE p=4e-13).

In summary, reduced network activity in TG neurons is contributed mainly by the deficits in neuronal architecture and to a lesser extent by over-inhibition, and EE effects on neuronal morphology are the main factor contributing to the improvement of network activity.

### Effects of EE on cognitive performance, levels of DYRK1A kinase activity, dendritic complexity and spine morphology are lost after EE discontinuation (EEdis) in TG mice

The rescue of recognition memory we detected upon EE in TGEE (2mo) mice was not maintained four months after discontinuation of EE (EEdis) in TG mice (TGEEdis; 6mo of age). TGEEdis (6mo) mice showed impaired object discrimination compared to TGEE (2mo) mice (Fig. 4a; three-way ANOVA discontinuation effect F(1,117)= 5.11, p=0.03; TGEEdis (6mo) vs. TGEE (2mo) p=0.06). Since our EE conditions did not improve recognition memory in WTEE (2mo) mice, EE discontinuation did not have an effect on WTEE mice when we evaluated novel object recognition test at 6 months of age (WTEEdis; 6mo) as they performed the novel object recognition test to a similar level than WTNE (6mo). The normalization of DYRK1A kinase activity achieved by EE was also lost in 6mo TG mice after EEdis, returning to NE increased activity levels (Fig. 4b; DYRK1A kinase activity: WTNE (6mo) = 1.00 +/− 0.12; TGNE (6mo) = 1.69 +/− 0.21; WTEEdis (6mo) = 1.04 +/− 0.24; TGEEdis (6mo) = 1.68 +/− 0.14). Neither object discrimination performance nor DYRK1A kinase activity results were caused by any age-related effect (Fig. S9a and S9b; three-way ANOVA age effect N.S.).

**Figure 4.**
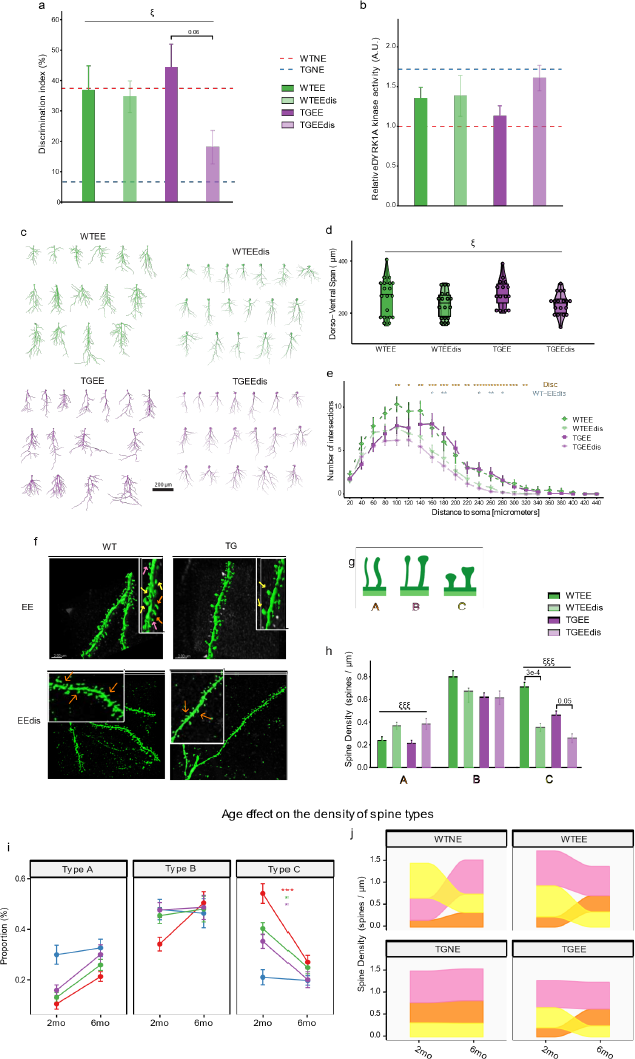
Long term effects on cognitive performance, levels of DYRK1A kinase activity, dendritic complexity and spine morphology are lost after environmental enrichment discontinuation in TgDyrk1A mice. **a** Discrimination index of the NOR paradigm in WT and TG mice reared on EE and reared on EE followed by four months under NE conditions (EEdis): WTEE (2mo) = 16; TGEE (2mo) = 19, WTEEdis (6mo) = 18; TGEEdis (6mo) = 18. Linear mixed effects, treatment discontinuation effect ξ p<0.05. **b** Relative DYRK1A kinase activity in the hippocampus. WTEE (2mo) = 12; TGEE (2mo) = 11; WTEEdis (6mo) = 12; TGEEdis (6mo) = 12. Data are expressed as mean ± SEM. Three-way ANOVA. Colored dashed lines indicate reference mean values for 2mo WTNE and TGNE groups. **c** 2D projection of all reconstructed apical dendritic trees (coronal plane) in WT and TG, upon EE (2mo) and after EEdis (6mo). Scale bar = 200 μm. **d** Length of CA1 apical dendritic trees along the dorso-ventral axis. Data are presented as an overlay of dotplots, boxplots and violin plots. WTEE (2mo) = 19/4, TGEE (2mo) =15/3; WTEEdis (6mo) = 18/3; TGEEdis (6mo) = 18/4. Linear mixed effects, treatment discontinuation effect ξ p<0.05. **e** Sholl analysis. Lines indicate mean ± SEM. Linear mixed effects, * p<0.05, ** p<0.01. **f** Representative confocal images showing CA1 apical dendritic spines in WT and TG mice under EE (2mo) or discontinued EE (6mo) conditions. Scale bar = 200 μm. Right upper insert corresponds to a higher magnification. Orange, pink and white arrows denote type A, type B and type C spines, respectively (**g**). **g** Illustration of the morphological categories used to classify spines. **h** Density of each spine type. Data are presented as mean ± SEM. Linear mixed effects, treatment discontinuation effect ξξξ p<0.001. **i** Age-related changes in the percentage of spine types (A, B, C). Data are presented as mean ± SEM. Data are paired along time for each experimental group (colored lines). Linear mixed effects, * p<0.05, *** p<0.001. **j** Sankey plots of the density of each spine type for WT and TG mice both NE and after EE treatment (2 and 6 mo) and its discontinuation (6mo). Orange, pink and yellow represent spines of types A, B and C, respectively. The width of each band indicates spine density in number of spines per micrometer. (f-j) Number of dendrite segments/neurons reconstructed/animals analyzed: WTNE-2mo 19/19/4; TGNE-2mo 16/12/4; WTEE (2mo) = 19/19/4, TGEE (2mo) = 16/9/4; WTNE-6mo 20/20/4; TGNE-6mo 19/19/5; WTEEdis (6mo) = 18/18/3; TGEEdis (6mo) = 18/18/6).

The width of apical CA1 was smaller in both 2mo and 6mo TGNE mice compared to WTNE (Figs. S3a and S3b; linear mixed effects genotype effect for NE groups F(1,31.88)=17.03, p=3e-4; TGNE-2mo vs WTNE-2mo p=0.02), due to a specific reduction of *SR* width (Fig. S3c; F(1,31.08)=31.75, linear mixed effects genotype effect p=4e-6; TGNE-2mo vs WTNE-2mo p=0.003; TGNE-6mo vs WTNE-6mo p=0.04). The rescuing effect of EE on the width of CA1 *SR* in TGEE mice was no longer significant upon discontinuation (Fig. S3a-c; linear mixed effects genotype x treatment interaction for EEdis (6mo) groups N.S., age effect N.S., genotype effect for EEdis (6 mo) groups F(1,38.20)=18.21 p=1e-4). Dendritic complexity was also reduced to NE levels in the Sholl analysis in both WT and TG mice (Fig. 4c and 4e; treatment discontinuation effect p<0.05 between 100 and 320μm and WTEEdis (6mo) vs WTEE (2mo) p<0.05 between 160 and 180, and 240 and 280μm. Fig. S4b; linear mixed effects, discontinuation effect F(1,62.84)=21.36, p=2e-5; TGEEdis (6mo) vs TGEE (2mo) p=3e-3), as was the dorsoventral span of apical dendrites (Fig. 4d; linear mixed effects discontinuation effect F(1,49.44)=6.64, p=0.01; TGEEdis (6mo) vs. TGNE (6mo), N.S.). To discard a possible influence of age in the effects of EE discontinuation, we compared 6 months old WTNE and TGNE (WTNE-6mo and TGNE-6mo) to their younger 2mo counterparts (WTNE-2mo and TGNE-2mo). We found no significant age effects in neither WTNE nor TGNE mice for *SR* width, dorso-ventral span or Sholl AUC (Fig. S3c, S4a and S4c linear mixed effects age effect N.S.). These results suggest that the dendritic remodeling effect of EE in TG hippocampal CA1 is completely lost upon EEdis.

Finally, as occurred with the other measures, discontinuation of EE also led to the loss of the treatment effect on spine density in TGEEdis mice, with a decrease in mature (type C) spines (Fig. 4f-h; Type C linear mixed effects discontinuation effect F(1,30.44)=37.29, p=1e-6; TGEEdis vs TGEE, p=0.05). These results are not age-dependent, since all spine classes remained unchanged with age in TGNE mice (Supplementary Figs. 4i and 4j). Instead, we found a significant reduction of mushroom-like type C spines in 6mo WT mice compared to 2mo WT (Figs. 4m and 4n; Type C WTNE-2mo vs. WTNE-6mo, p=6e-8) that may explain that this time we detect an effect of EEdis in WT mice (Fig. 4l and 5m; Type C WTEEdis vs WTEE, p=3e-4). No age-related differences in the total density of dendritic spines of CA1 pyramidal neurons located in the *SR* layer were observed in non-enriched groups (Fig. S4c).

**Figure 5.**
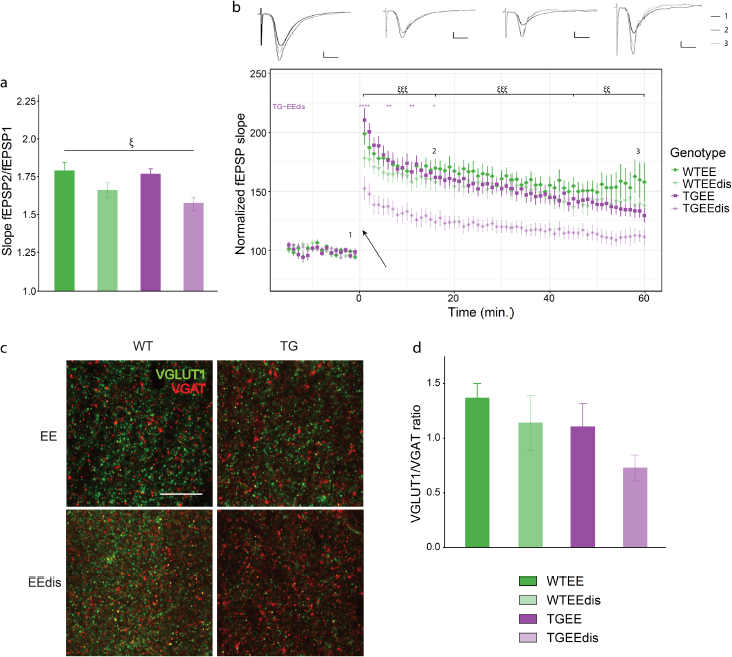
Changes in synaptic plasticity are lost in TgDyrk1A mice after EE discontinuation. **a** Paired pulse facilitation (fEPSP2/fEPSP1) in the hippocampus with an inter-stimulus interval of 50ms of WT and TG mice reared under EE conditions for one month (2mo) or after environmental enrichment discontinuation (EEdis, 6mo). Slices/animals: WTEE (2mo) = 9/5; TGEE (2mo) = 16/5; WTEEdis (6mo) = 20/7; TGEEdis (6mo) = 12/5. Three-way ANOVA treatment discontinuation effect ξ p<0.05. **b** Long-term potentiation (LTP) in CA1 induced by theta burst stimulation (TBS) in CA3. Upper panel: representation of the mean of 5 responses to stimulation at time 1 (−5 min), 2 (15 min) and 3 (60 min). Vertical bar represents 500µV, horizontal bar represents 5ms. Lower panel: normalized fEPSP slope before and after TBS. Black arrow represents the time of TBS application (t=0). Three-way ANOVA; ξξ p<0.01, ξξξ p<0.001. **c** Representative confocal images showing VGLUT1+ and VGAT+ punctae in the CA1 *SR* hippocampal subfield. Scale bar = 10 μm. **d** Histogram showing the ratio between VGLUT1+ and VGAT+ punctae. WTEE (2mo) n = 8; TGEE (2mo) n = 7; WTEEdis (6mo) n = 5; TGEEdis (6mo) n = 4. Data are represented as mean + SEM.

### EE-induced improvement of synaptic plasticity is lost upon EE discontinuation in TgDyrk1A mice

Upon discontinuation, the EE effects on paired-pulse facilitation (Fig. 5a, three-way ANOVA discontinuation effect F(1,105)=8.92, p=0.004), and on LTP were lost in TGEE-6mo (Fig. 5b; three-way ANOVA, discontinuation effect p<0.05; TGEEdis (6mo) vs TGEE (2mo), p<0.05). This was not due to an age-effect on the LTP response (Fig. S10a; three-way ANOVA age effect N.S.; WTNE-2mo vs WTNE-6mo and TGNE-2mo vs TGNE-6mo N.S.). We detected a significant genotype-related reduction in LTP for both 2mo and 6mo TGNE mice (Fig. S10a; three-way ANOVA genotype effect for NE groups F(1,105)=5.68, p=0.02). In addition, the excitation-inhibition imbalance showed a significant genotype effect in NE groups at both 2mo and 6mo mice (Fig. S10b; three-way ANOVA genotype effect for NE groups F(1,41)=7.57, p=0.009), showing a non-significant trend to decreased VGLUT/VGAT ratio after EEdis (Fig. S10b; three-way ANOVA discontinuation effect F(1,41)=3.63, p=0.06). Those results were also not dependent on age.

### EE effects on the connectivity repertoire and input-output frequency relationship were lost after treatment discontinuation in both TG and WT mice

When only accounting for dendritic tree span and complexity, the increase of connectivity repertoire detected with EE in WT and TG was lost upon EE discontinuation (Fig. S5a; linear mixed effects discontinuation effect F(1,57.49)= 18.706, p=6.14e-5, TGEE (2mo) vs. TGEEdis (6mo) p=0.003). Similarly, when considering mature spine densities, the effect of EE was lost (Fig. S5b; linear mixed effects discontinuation effect F(1,61.35)=41.34 p=2e-8, WTEE (2mo) vs. WTEEdis (6mo) p=0.001 and TGEE (2mo) vs. TGEEdis (6mo) p=2e-4), so that both WTEEdis (6mo) and TGEEdis (6mo) neurons showed the same levels of connectivity repertoire as NE 6mo counterparts (Fig. S5b). As such, the genotype-dependent reduction of connectivity repertoire found in TGNE mice at 2mo was also present at 6mo of age (genotype effect for NE groups F(1,33.65)= 25.29 p=2e-5; WTNE-2mo vs. TGNE-2mo p=0.02 and WTNE-6mo vs. TGNE-6mo p=0.001). Connectivity repertoire did not significantly change with age in either genotype in NE groups (Fig. S5b).

Upon in silico LF and HF stimulation of the whole tree, the EEdis (6mo) neurons showed a significant reduction in their success spiking rate and firing rate compared to enriched ones EE (2mo) in both genotypes (Fig. S8a; linear mixed effects discontinuation effect F(1,53.85)= 35.24, p=2e-7; TGEEdis (6mo) vs. TGEE (2mo) p=4e-7; Figs. S11c and S8b; linear mixed effects discontinuation effect F(1,51.28)= 56.85 p=7e-10; WTEEdis (6mo) vs. WTEE (2mo) p=4e-4; TGEEdis (6mo) vs. TGEE (2mo) p=1e-5). Neither WTEEdis nor TGEEdis neurons were able to reach high γ (Fig. 11e). None of these effects could be attributed to age-related differences in success rate, or the output firing as neither WTNE-6mo nor TGNE-6mo showed no significant differences to WTNE-2mo and TGNE-2mo in any of them (WTNE-6mo reached high γ as WTNE-2mo and TGNE-6mo did not, as TGNE-2mo; Fig. S7a and S7b). When simulating inputs on only *SR*, both WT and TG neurons had a slight reduction in the output firing rate with EE discontinuation (See <u>Supplementary Text</u> and figure S11).

In summary, our results suggest that after discontinuation, the effect of EE on the response to HF inputs in the whole dendritic tree was lost in TGEEdis neurons, which were still unable to reach high γ frequencies. This is due to the loss of EE effects on dendritic tree complexity, that, along with decreased mature spine density leads to reduced coincident input integration in both WTEEdis and TGEEdis neurons.

### EE effects on the activity of CA1 neuronal populations are lost after treatment discontinuation

When simulating HF stimulation on the CA1 neuronal population this time accounting also for the inhibition phenotype, the activity of the network was markedly reduced in WTEEdis (6mo) and TGEEdis (6mo) compared to enriched conditions (2mo) (Fig. 6a; 72% reduction in the power Figs. 6b and 6c three-way ANOVA discontinuation effect F(1,32)= 637.17, p=1e-22; WTEEdis (6mo) vs WTEE (2mo), p=6e-21; TGEEdis (6mo) vs TGEE (2mo), p=3e-11). The effect of EE discontinuation is partly explained by a significant age-dependent reduction in the network activity we found in WT (14%) and TG (39%) (Fig. 6c; thee-way ANOVA age effect for NE groups F(1,32)= 18.59, p=2e-4; TGNE_6mo vs TGNE_2mo, p=0.02; WTNE_6mo vs WTNE_2mo, p=0.02). Additionally, we found a deleterious effect of EE discontinuation, as WTEEdis showed reduced neuron population activity in comparison to WTNE_6mo (42%; WTNE_6mo vs WTEEdis, p=1e-6).

**Figure 6.**
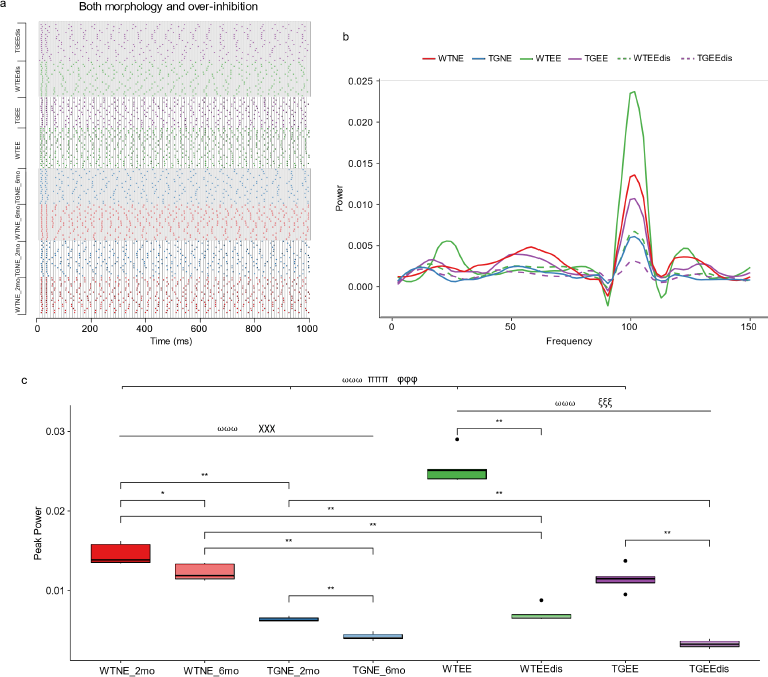
EE effects on averaged activity of TgDyrk1A CA1 neurons are lost after treatment discontinuation. **a** Spike raster plot of the simulated activity of CA1 neurons of WTNE (red), TGNE (blue) WTEE (green) and TGEE (purple), taking into account both. Note the reduced firing in both WT and TG NE_6mo neurons upon 100 Hz stimulation that is more marked upon Eedis in WTEEdis and TGEEdis. Raster plots overlayed by gray blocks indicate 6mo and EEdis groups. **b** Power spectra of spiking activity obtained upon simulation of 100 Hz input frequency and averaged over 10 simulation repetitions for WT and TG NE, EE and EEdis groups. Dashed lines show the results of simulations accounting for EEdis. **e** Peak power at 100Hz for each of the neuron populations presented as boxplots. Number of neurons/animals: WTNE = 18/5; TGNE = 18/5; WTEE = 18/3; TGEE= 18/4. Three-way ANOVA, genotype effect ⍵⍵⍵ p<0.001; treatment effect πππ p<0.001; genotype x treatment interaction φφφ p<0.001; treatment discontinuation effect ξξξ p<0.001; age effect χχχ p<0.001. * p<0.05, ** p<0.01.

## DISCUSSION

It has long been suggested that dendritic abnormalities and synaptic plasticity alterations are landmarks of intellectual disability, and especially of DS^19^. This concept has driven therapeutic approaches focused in boosting neural plasticity, with the goal that their correction would contribute to cognitive improvement. Such correction has been shown with environmental intervention (EE) interventions that partially rescued microstructural phenotypes and cognition^8,9,11,20^. We here used a mouse model overexpressing *Dyrk1A*, one of the most relevant DS candidate genes to understand the functional implications of dendritic alterations and their rescue by EE.

Our behavioral experiments recapitulated the impaired performance in the Novel Object Recognition^21^. TG mice showed significantly reduced object discrimination, as described in other DS^22^ and transgenic *Dyrk1A* models^23^. We also identified structural and functional hippocampal alterations in TG mice similar to DS hippocampal phenotypes^8,9,11,20,24^, and an imbalance between excitatory and inhibitory neurotransmission, biased towards inhibition, mainly due to a reduced number of VGLUT1 puncta (excitatory vesicles) ^25 26^, along with impaired LTP in CA3-CA1 and differences in structural organization of CA1 subfield. Specifically, we observed decreased apical dendritic tree span and complexity, and a more immature spine phenotype in TG pyramidal CA1 neurons. Dendritic branching alterations and reduced mature spine density in TG CA1 neurons reduce the number of afferent inputs and the possible input combinations, as shown by our connectivity repertoire analysis. This is relevant as dendritic arbor architecture and span play key roles in local computation processes^27–31^, such as integration of pre- and post-synaptic events.

An interesting observation of our work is that the dendritic alterations were layer-specific, with reduced dendritic width affecting mainly the *stratum radiatum* in the TG CA1 subfield. Thus, the next important question was how to integrate these data to infer network alterations in TG CA1. To this aim we developed a computational model in which we could integrate those deficits and model the possible outcomes at the single cell and network levels. First, as a reduced connectivity repertoire may affect neuronal signal integration^16^, we explored the input-output (IO) relation of firing frequencies in individual CA1 pyramidal neurons using multi-compartmental models. This allowed simulations of either the complete tree, spanning *SR* and *SL,* or of the dendritic area covering *SR* alone selectively, as the reduced width was mainly dependent on *SR*. In WT neurons it is expected that SC stimulation only in *SR* (even with very high-frequency stimulation) will produce selective output firing at low γ^17^, a property called frequency selectivity. Instead, combined input from SC and the perforant path to both *SR* and *SL* can lead to higher frequency output firing through coincidence detection^32,33^. This is relevant because frequency selectivity allows CA1 cells to selectively synchronize with CA3 (at low γ) or the entorhinal cortex (at high γ), during familiar environment exploration or novel object-place pair investigation^34,35^. In fact, it has been shown that prospective coding occurs when CA1 cells mainly fire at low γ during spatial exploration tasks^36^. To assess whether frequency selectivity (e.g. the fact that CA1 to selectively “listen” to CA3 at slow compared with fast gamma) could be impaired in TG CA1 neurons (taking into account the reduced synaptic connectivity resulting from the dendritic alterations we detected), we simulated receiving only SC inputs (e.g. affecting only *SR*) or in synapses receiving SC and perforant path inputs (*SR* and *SL*). With only *SR* inputs in the model, TG neurons were not able to reach low γ. When we stimulated *SR* and *SL*, TG neurons were not able to reach high γ. This suggests that the ability to switch to high γ coupling between CA1 and EC could be impaired in TG mice. Supporting this hypothesis, Munn et al. have showed that Ts65Dn CA1 pyramidal cells are more likely to be phase-locked to slow than fast gamma^37^. Thus, a perturbed γ coupling could impair the ability to distinguish familiar from novel environments^34,35^.

The reduced ability of single neurons to reach high γ suggests that architectural features are relevant to produce alterations of rhythmic activity at the local neuronal population level. In TgDyrk1A and trisomic models, neuronal population activity is altered in cortical and hippocampal local networks, with decreased high γ activity^5,38^, an effect that has been mainly attributed to excessive GABA-mediated neurotransmission in DS^39,40^. We tested *in silico* the impact of morphological alterations on the ability of a simulated CA1 neuronal population formed by the neurons we traced to fire synchronously at high γ. To do so, we stimulated the multicompartmental models of the neurons with simultaneous input and we simulated recurrent local inhibitory feedback including in the model point inhibitory neurons that both inhibit pyramidal neurons and themselves. With these simulations, we showed that reduced spine density, and dendritic tree span and complexity, have a stronger impact than over-inhibition from local circuits on CA1 neurons. These results indicate that the excess of GABAergic transmission is only a partial explanation for the alterations in population activity rhythms, and that architectural features should also be considered.

EE rescued the rescued recognition memory, dendritic abnormalities, and synaptic plasticity alterations in TG mice, as was also reported for other DS mouse models and in humans^8,9,11,20^. EE has been shown to increase spine density and to stabilize new synapses^41^. In TG mice, EE increased mature spine density, suggesting that it stimulated the maturation and stabilization of preexisting spines, but did not promote spinogenesis, similar to previous findings^42–44^. Along with the significant increase in mature spine density, EE produced a non-significant increase in dendritic span and complexity, that together were sufficient to improve the connectivity repertoire. The treatment also led to a significant increase of the VGLUT/VGAT ratio. Nevertheless, our simulations show that only correcting the excitation-inhibition balance would not be sufficient to rescue fast γ. This is relevant given that previous work^45^ showed that the reduction of inhibition in Ts65Dn with the GABA_A_ antagonist pentylenetetrazol in the dentate gyrus rescued cognitive impairment. Instead GABAA α5 subunit inverse agonist treatment did not meet its primary and secondary endpoints on improvement in cognition in a recent clinical trial (RG1662; Hoffman-La Roche). Given that our results suggest that only a treatment rescuing both dendritic architecture and GABAergic synaptic signaling levels would rescue the network alterations in TG mice, it might be speculated that administration of a drug which besides reducing inhibition would also increase mature spine density and dendritic morphology as has been reported for pentylenetetrazol in rat CA1 neurons^46^ and snails^47^ would be more effective that a treatment which decreases dendritic arborization^48^ and spine maturation^49^ as reported for GABA_A_ α5 subunit inverse agonist treatment in hippocampal neurons *in vitro*. Thus, further work to elucidate the impact of GABA_A_ antagonists on dendritic architecture is needed to explore improved pharmacotherapy for DS.

Another main novel aspect of this work is the analysis of the stability of EE effects upon discontinuation. The main concern of early intervention in DS is that cognitive improvements are not maintained after treatment discontinuation, thus demanding costly life-long interventions to retain the positive effects^20,50^. To understand if the overexpression of *Dyrk1A* could be involved in the instability of this phenotypic rescue, we studied previously EE exposed animals after four months from EE discontinuation. Under these conditions, the rescuing effects of EE on dendritic structure, spine morphology hippocampal CA1 synaptic plasticity and short-term memory, were lost in TG mice. Importantly, we observed that Dyrk1A kinase activity, normalized by one month of EE^10^, returned to non-enriched levels in TG mice. Taken together, these results suggest that *Dyrk1A* overexpression impedes the maintenance of activity-dependent neuronal plasticity improvements in DS pathology. When introducing these observations in our computational model, we observed that after EE discontinuation TG neurons showed firing frequencies comparable to their NE counterparts both at the single cell and local population levels. While the effects of EE discontinuation in WT did not have a significant impact on object discrimination nor DYRK1A kinase activity, they showed a significant decrease in mature spine density and a non-significant decrease in VGLUT/VGAT puncta ratio. The combination of those led to a significantly decreased γ firing at the local population level. While LTP was not impaired in WT after EE discontinuation, further exploration is needed to corroborate such putative impairment and the associated spine phenotype.

In conclusion, our experiments showed that behavioral impairment in novel object recognition is accompanied by structural alterations at the cellular and subcellular scale and altered LTP electrophysiology. Those were temporarily rescued by EE, which implied remodeling of distal dendrites in CA1 pyramidal cells and a recovery of both mature spines and excitation-inhibition balance. We used a computational model to understand the link between the structural perturbations and neuronal function, showing that reduced inputs impair the ability of single neurons to reach high frequency firing. We conclude that synaptic and structural neuronal defects in TG mice lead to impaired cognition by impairing hippocampal CA1 integration of CA3 and entorhinal cortex (EC) inputs and that excessive inhibition is only partially responsible for decreased local population oscillatory activity, suggesting that microarchitectural neuronal features are more relevant than previously thought.

## Supporting information

Resource Table

## ACKNOWLEDGMENTS

We acknowledge support of the Spanish Ministry of Economy and Competitiveness (MINECO), ‘Centro de Excelencia Severo Ochoa 2013-2017’, SEV-2012-0208. The research leading to these results has received funding from MINECO (SAF2013-49129-C2-1-R), the NIH grant R01 EB028159 and Lejeune Foundation. All confocal imaging was done in the CRG Advanced Light Microscopy Facility. We acknowledge the support from Gemma Comas for their help imaging the single neuron slice sections. We are grateful to Santiago Acosta for his help in generating the single neuron reconstructions, and Juan Luis Musoles for his help in the preprocessing of CA1 lower magnification image stacks.

## AUTHOR CONTRIBUTIONS

Conceptualization, M.D. and M.M.L.; investigation, M.PE, L.MG., T.G., I.BY.; project administration, M.D.; writing – original draft, M.PE., L.MG., M.M.L. and M.D.; writing – review & editing, L.MG., M.P., T.G., I.BY., M.M.L. and M.D.

## KEY RESOURCES TABLE

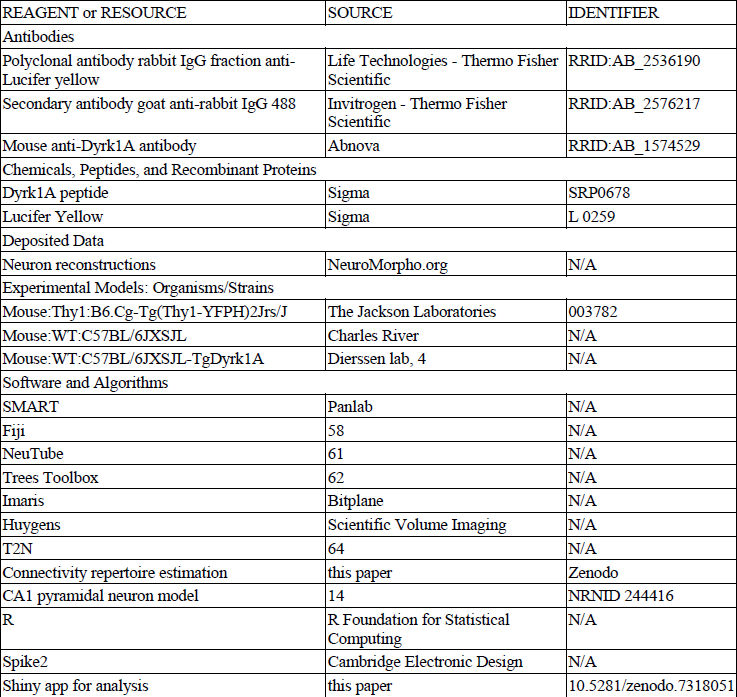

## RESOURCE AVAILABILITY

### Lead contact

Further information and requests for resources and reagents should be directed to and will be fulfilled by the lead contact Prof. Mara Dierssen –.

### Materials availability

This study did not generate new unique reagents.

### Data and code availability

Neuron reconstructions have been deposited at the NeuroMorpho.Org repository and at GitHub: https://github.com/lmanubens/EE_CA1/tree/main/data. Source data for all the analyses can be found at: https://github.com/lmanubens/EE_CA1/tree/main/shiny_app.

All original code for the Shiny app, statistical analysis and plotting has been deposited at GitHub and is publicly available as of the date of publication: https://github.com/lmanubens/EE_CA1/tree/main/shiny_app. Code for multi-compartmental modeling can be found at: https://github.com/lmanubens/EE_CA1/tree/main/model. Any additional information required to reanalyze the data reported in this work paper is available from the corresponding authors upon request.

## METHODS

### Experimental model and subject details

#### Animals

The production of mice transgenic for *Dyrk1A* (TgDyrk1A) has previously been described^4^. The transgene was inserted into C57BL/6JXSJL (Charles River, Barcelona, Spain) embryos and the stock is maintained by intercrossing wild type and TgDyrk1A mice derived from the original founder. The non-transgenic littermates served as controls. For the experiments to measure the width of the hippocampal layers, we generated double transgenic mice (Thy1-YFP/TgDyrk1A) by crossing TgDyrk1A^4^ male mice with Thy1-Yellow Fluorescent Protein (YFP) females (strain B6.Cg-Tg(Thy1-YFPH)2Jrs/J n°003782; The Jackson Laboratories). Same sex littermates were group-housed under a 12-h light/dark schedule (lights on at 09:00 a.m.) in controlled environmental conditions of humidity (60%) and temperature (22 ± 2 °C) with free access to food and water. The present experiments were conducted using only females, since TG male mice showed hierarchical behavior (data not shown), similar to that observed in other DS mouse models^8^ that may affect the outcome of EE. Even so, our non-enriched females showed the same phenotypes as we previously detected in TG males^10^. The non-transgenic littermates served as controls. For all the analyses we compared WT and TG mice reared in NE conditions, after one month of EE conditions and after EE discontinuation during four months. Given that EE effects after discontinuation could be confounded with age-associated changes between 2- and 6-months mice, we also assessed the age effect separately.

All experiments followed the principle of the “Three Rs”: replacement, reduction and refinement according to Directive 63 / 2010 and its implementation in Member States. The study was conducted according to the guidelines of the local (law 32/2007) and European regulations (2010/63/EU) and the Standards for Use of Laboratory Animals no. A5388-01 (NIH), and approved by the Ethics Committee of Parc de Recerca Biomèdica (Comité Ético de Experimentación Animal del PRBB (CEEA-PRBB); MDS 0035P2). Reporting followed the ARRIVE (Animal Research: Reporting of In Vivo Experiments) guidelines with the modifications suggested by the Trisomy 21 Research Society for work with DS mouse models. The CRG is authorized to work with genetically modified organisms (A/ES/05/I-13 and A/ES/05/14).

#### Housing and enrichment conditions

After weaning (21 days of age), TG and WT mice were randomly reared under either non-enriched (NE) or enriched (environmental enrichment; EE) conditions for 30 days (Supplementary Fig. 1a). For the NE conditions, animals were reared in conventional Plexiglas cages (20 x 12 x 12 cm height) in groups of two to three animals. The EE group was reared in spacious (55 x 80 x 50 cm height) Plexiglas cages with toys, small houses, tunnels, and platforms, but without wheels to avoid the possible effects of physical exercise. The arrangement was changed every 3 days to maintain the novelty of the environment. To stimulate social interactions, six to eight mice were housed in each EE cage. All groups of animals were maintained under the same 12 h (8:00 to 20:00) light-dark cycle in controlled environmental conditions of humidity (60%) and temperature (22 ± 1°), with free access to food and water. To analyze the long-term stability of EE effects, after 1 month housed in EE conditions, a separate group of WTEE (WTEE discontinued; WTEEdis) and TGEE (TGEE discontinued; TGEEdis) mice was reared for 4 months under NE conditions (Supplementary Fig. 1b).

### Method details

#### Novel object recognition test

The novel object recognition test is widely used to assess recognition memory in rodents^51^. Briefly, mice were placed into an open-field (70 cm wide x 70 cm long x 30 cm high) made of black methacrylate, surrounded by curtains. The task was performed under non-aversive low lighting conditions (50 lux). An overhead camera connected to video-tracking software (SMART, Panlab) was used to monitor the animal’s behavior. To eliminate odor cues, the arena and the objects were thoroughly cleaned with 10% odorless soap, and dried. In addition, the positions of the objects in the familiarization and test session were counterbalanced between animals. Sniffing of the objects was used as a measure of exploration and was registered manually by an experimenter blind to genotype and treatment. The first day, mice were allowed to explore the arena for 5 min for familiarization with the experimental setting. Thereafter, an object was placed in the centre of the arena, and mice were given the chance to habituate. Exploration of the object was recorded for 5 min. On the second day (Supplementary Fig. 1c), during the familiarization trial, mice were allowed to explore two identical objects placed in the center of the apparatus for 10 min. Mice exploring less than 20 s were discarded. One hour after, during the test session, mice were allowed to explore the same arena for 5 min, but with one of the familiar objects exchanged by a new one. The exploration times for the familiar (TF) and new object (TN) during the test phase were recorded (number of animals: WTNE = 14; TGNE = 15; WTEE = 16; TGEE = 19 and after EE discontinuation: WTNE = 11; TGNE = 14; WTEEdis = 18; TGEEdis = 18). Memory was operationally defined as the discrimination index (DI), defined as the time spent investigating the novel object minus the time spent investigating the familiar one during the testing period [Discrimination Index, DI = [(TN – TF) / Total Exploration Time] x 100].

#### Neuromorphological analysis

The structural changes were analyzed on the dorsal hippocampal CA1 layer^52^ (Mouse Brain Atlas: antero-posterior = −1.34 to −2.54 mm; medio-lateral 0.5 to 1.6 mm; dorso-ventral 1.2 to 2 mm; Supplementary Video 1) of Thy1-YFP/TgDyrk1A (TG) and Thy1-YFP/WT. Data from the left and right hemispheres were taken indistinctly.

##### Layer width analysis of hippocampal CA1 strata

Mice were sacrificed and perfused intracardially with phosphate buffered saline (PBS), followed by chilled 4% paraformaldehyde (PFA; Sigma). The brains were removed from the skull, postfixed in the same fixative at 4°C overnight, and cryoprotected in 30% sucrose. One-hundred fifty μm coronal brain sections were obtained using a vibratome (VT1000S, Leica Microsystems), washed extensively with 0.1M PBS, and mounted and coverslipped with mowiol (number of slices/animals: WTNE = 14/5; TGNE = 8/4; WTEE = 17/6; TGEE= 8/3) and 4 months after termination of EE treatment to evaluate the long-term stability of EE effects (number of slices/animals: WTNE = 24/8; TGNE = 9/4; WTEEdis = 18/6; TGEEdis = 19/7).

Images of the dorsal hippocampal CA1 region were obtained using a confocal microscope (SPE; Leica Microsystems) with a 10x objective. A line scan of 1024 × 1024 pixels, 3µm wide z steps and 3 frame intensity averages were used. An average projection was generated with the Fiji imaging analysis software (version 2.9.0/1.53t)^53^. Image stacks of slanted vibratome sections were discarded for subsequent analysis.

The width of the *stratum radiatum* (intercept between the *stratum pyramidale-radiatum* border and the *radiatum-lacunosum* border) and *stratum lacunosum* (intercept between the *stratum radiatum-lacunosum* border and the hippocampus sulcus), where apical dendrites of CA1 pyramidal neurons are located, were analyzed at four medio-lateral locations^52^ (Bregma, antero-posterior = −1.06 to −2.54 mm; medio-lateral 0.5 to 1.6 mm) using the Fiji straight line selection tool. The four locations were equispaced by ∼200µm from proximal to distal along the medio-lateral axis (Fig. S2).

##### Single-cell morphological analysis of dendritic tree architecture of hippocampal pyramidal cells

To reconstruct dendritic tree architecture of single CA1 pyramidal cells, TG animals were perfused with 4% PFA, and 150 μm coronal sections from the dorsal hippocampal CA1 region (Bregma, antero-posterior = −1.06 to −2.54 mm; medio-lateral 0.5 to 1.6 mm) were obtained with a vibratome. Intracellular injections were performed in CA1 pyramidal neurons by continuous current of fluorescent Lucifer Yellow (LY) as described in detail in Elston et al.^54,55^. Briefly, pyramidal neurons located randomly throughout the dorsal CA1 area^52^ (Bregma, antero-posterior = −1.06 to −2.54 mm; medio-lateral 0.5 to 1.6 mm) were selected for LY injection and further reconstruction (number of neurons/animals: WTNE = 18/4; TGNE = 19/4; WTEE = 19/4; TGEE= 15/3 and WTNE = 18/5; TGNE = 18/5; WTEEdis = 18/3; TGEEdis= 18/4). With a preliminary set of reconstructions (N=6 for WTNE and TGNE) we did a sample size estimation based on the total length measured in those neurons (mL_WT_ =1637, mL_TG_ =1066, SD=536). A sample size of N=14 would have a Type I error rate of 0.05 and a power of 0.80. The sections were then counter-stained with antibodies against LY. Sections were washed with PBS and 0.3% PBS-T to make the cells permeable. To minimise the background staining, the slices were treated with 50 mM glycine (minimum 99% TLC, Sigma-Aldrich, St. Louis, MO, USA) in 0.3% PBS-T for 20 min. Thereafter, samples were treated for 2 h at room temperature with 3% bovine serum albumin (BSA, Sigma-Aldrich, St. Louis, MO, USA), and 0.3% PBS-T as a blocking agent, and were incubated overnight at 4°C in polyclonal rabbit IgG fraction anti-Lucifer yellow (1:500, Life Technologies - Thermo Fisher Scientific, RRID:AB_2536190) in 0.3% PBS-T and 1% BSA. The slices were then incubated 2h at room temperature in goat anti-rabbit IgG 488 (1:200, Invitrogen - Thermo Fisher Scientific, RRID:AB_2576217, Carlsbad, California, USA) in 1% BSA in 0.3% PBS-T. Finally, sections were washed with 0.3% PBS-T and PBS and coverslipped with mowiol mounting medium.

Image analysis and neuronal reconstruction: Images of single neuron apical trees for neuronal reconstruction were acquired with a confocal microscope (SP5 Upright; Leica Microsystems) with a 20x air objective (HCX PL APO CS 20.0×0.70 dry UV). A line scan of 1024 × 1024 pixels, 0.347µm wide z steps and 3-line intensity averages were used for imaging whole dendritic trees. Leica Smart Gain was used to equalize the fluorescence signal intensity in depth in the samples. The signal to noise ratio was measured for each stack at a 100µm depth for a representative dendritic segment and an equivalent area containing only background at the same depth. Images with SNR<1.5 were discarded for subsequent analysis. 2D gaussian blurring was applied to the stacks with a sigma of 2 pixels https://imagej.nih.gov/ij/developer/api/ij/plugin/filter/GaussianBlur.html, background subtraction (50 pixel sliding paraboloid without smoothing; https://imagej.net/Rolling_Ball_Background_Subtraction) and contrast enhancement (with 0.3% saturated pixels and using the stack histogram; https://imagej.nih.gov/ij/developer/api/ij/plugin/ContrastEnhancer.html) in Fiji.

Dendritic trees were reconstructed with NeuTube (version 1.0z.2018.07)^56^. Morphological metric statistics were obtained with the TREES Toolbox (version 1.15) function stats_tree, documented in detail at https://www.treestoolbox.org/manual/stats_tree.html^57^. Briefly, the function provides a set of measurements that describe in detail morphological properties of the dendritic tree: total length, maximum and mean path length, mean branch length, dendritic spanning volume, number of branch points, maximum and mean branching order, dendritic area (surface occupied by dendrite), mean straightness of dendritic branches, mean branching angle, mean asymmetry, center of mass in different directions and the ratio between width and height (see Supplementary Table 1 for details). We measured the extent of the reconstructed neurons within CA1 *stratum radiatum* and *lacunosum* by aligning the apical tree in the dorso-ventral direction. Additionally, the stats_tree function allows us to obtain the Sholl analysis for each of the trees. For our analyses we used 20µm separation between consequent Sholl spheres at which the number of intersections were quantified. The data was plotted using the ggline of the R (version 4.1.0) package ggpubr (version 0.4.0).

##### Dendritic spine density and morphology

Images from the LY-injected CA1 pyramidal neurons used to analyze dendritic spine density and morphology from single neuron reconstructions were acquired with a confocal microscope (SPE; Leica Microsystems) using a 63x oil objective (ACS APO 63.0×1.30 oil) plus 5 times magnification. A line scan of 1024 × 1024 pixels, 0.1µm wide z steps and 3 frame intensity averages were used for imaging dendritic segments. Representative images were obtained by 3D rendering of the confocal imaging stacks in the Imaris software (Bitplane) after image deconvolution using Huygens (version 15.05; Scientific Volume Imaging). The resulting dendritic spine images were quantified with the aid of Fiji.

Dendritic spine density was measured by quantifying the number of spines in 20 μm length dendritic segments located 30 μm distal from the soma of LY-injected CA1 pyramidal cells in 60-70% of neurons reconstructed (number of dendrite segments/neurons reconstructed/animals analyzed: WTNE = 19/19/4; TGNE =16/12/4; WTEE = 19/19/4; TGEE = 16/9/4 and after EE treatment discontinuation: WTNE = 20/20/4; TGNE = 19/19/5; WTEEdis = 18/18/3; TGEEdis = 18/18/6). Spines were classified as filopodia (type A), immature (type B) or mushroom (type C) (Fig 2i).

##### Excitation-inhibition balance analysis

In a separate experiment, mice were sacrificed, and perfused intracardially as detailed above (number of animals analyzed: WTNE = 7; TGNE = 7; WTEE = 8; TGEE = 7 and after EE treatment discontinuation: WTNE = 5; TGNE = 6; WTEEdis = 5; TGEEdis = 4). Forty μm coronal sections were obtained using a cryostat (CM3050S, Leica Microsystems). Analysis was performed in one of every six sections (6-8 sections per animal), covering the dorsal hippocampus^52^ (Bregma, antero-posterior −1.06 to −2.54 mm; medio-lateral 0.5 to 1.6 mm). Free-floating brain sections were permeabilized with 0.3% Triton X-100 in PBS for 30 min at room temperature (RT), and blocked with 20% fetal bovine serum and 0.3% Triton X-100 in PBS for one hour at RT. Subsequently, sections were incubated overnight at 4°C with the primary mouse anti-vesicular glutamate transporter 1 (VGLUT1) antibody (1:200, Synaptic Systems, RRID:AB_887879) to analyze presynaptic excitatory punctae and the guinea pig anti-vesicular GABA transporter (VGAT) antibody (1:200, cytoplasmic domain, Synaptic Systems, RRID:AB_887871) to evaluate presynaptic inhibitory punctae. The following day, while protected from light, slices were incubated with the corresponding secondary fluorescent antibodies (1:1000, Alexa 488 RRID:AB_2535764 and 555 RRID:AB_141784, Invitrogen) for one hour at RT.

Four images per section (8 sections per animal) were captured in the *stratum radiatum* of CA1 hippocampal subfield using a confocal microscope with a 63x objective (ACS APO 63.0×1.30 oil) and 5x magnification (SPE; Leica Microsystems). High-resolution images (pixel size = 0.05×0.05μm) were obtained and imported into Fiji and the negative control background intensity of each image was subtracted from each channel. To identify immunopositive punctae, the preprocessed images were thresholded and binarized using the Threshold command in Default mode. The number of VGLUT1 and VGAT puncta within the *stratum radiatum* area in CA1 were counted with a particle analysis plug-in of ImageJ using a defined puncta size range.

#### Modeling the functional consequences of dendritic remodelling

##### Connectivity repertoire

To study putative functional implications of genotype and treatment-dependent changes in dendritic architecture and spine density, we analyzed the connectivity repertoire *S* of the reconstructed neurons, defined as the possible connectivity patterns between dendrites and surrounding axons^58^. Specifically, we explored whether the genotype and/or treatment-dependent differences in neuronal shapes affect the repertoire of possible connectivity patterns between dendrites and surrounding axons. Briefly, following^58^ we measured total dendritic length with the *stats_tree* function of the TREES toolbox and the arbor radius *R* with a self-developed MATLAB script that provided the root mean square distance between any 2 dendritic segments from the 3D apical pyramidal dendrites reconstructions described above:

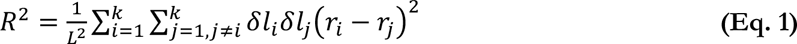

Being *L* the total length, *k* the total number of segments, *δl_i_* the length of the segment *i* and *r_i_* the position vector of the segment *i*. To obtain a quantification of the connectivity repertoire as a function not only of the dendritic tree morphology but also the density of synaptic contacts, which are strongly affected by the genotype and EE, we introduced spine densities measured in each group in the calculation of *S* by scaling the number of inputs in the tree *N* by the spine density *s_d_* resulting in the following equation:

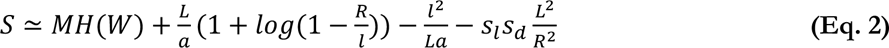

##### Input-output frequency relation

We also investigated whether the genotype and treatment dependent dendritic remodelling affect the efficiency of WT and TG trees to integrate synaptic signals. To this aim we explored the input-output frequency relationship defined as the evoked spiking frequency of a neuron upon the activation of 15% of available input synapses at different frequencies (from 0 to 100 Hz in steps of 10 Hz), since CA1 dendritic trees have been shown to lock at constant low gamma output frequency of 50Hz for high gamma input frequencies (ranging from 50 to 100Hz) at their *stratum radiatum*^17^. We based our model on Combe et al.^17^ (with ID in the ModelDB database NRNID 244416), in which the authors explore selective firing frequency of CA1 pyramidal neurons upon stimulation of Schaffer collateral input. They observed linear response for input frequencies between 0 and 40 Hz, and selective firing at slow gamma for input frequencies between 40 and 100Hz. Membrane channel conductances and distributions were mimicked from the template model. We modified the synaptic weight, following experimental measurements obtained in CA1 pyramidal neurons of the mouse^59^. We explored systematically the percentage of active synapses and found that 15% best recapitulate the behavior of the original model (spike-time adaptation as observed in electrophysiological experiment in mouse CA1) after updating the values of synaptic weights. In addition to the Schaffer collateral input to *stratum radiatum*, we also included perforant path synapses in *stratum lacunosum* in order to explore whether summation of both inputs would lead CA1 neurons to modify their firing frequencies to high gamma^33^.

To model this behavior computationally, we used the NEURON simulation environment (version 7.7.1) and T2N (version 1.13.1)^60^ in MATLAB (R2021a). T2N is an extension of the TREES toolbox, that allows to define passive and active membrane electrophysiological properties (i.e., passive currents, active ion channels, synaptic inputs, etc.) depending on the position within a neuronal reconstruction and export a multi-compartmental model in NEURON format, to simulate neuron dynamics upon injection of currents. In our case, as we were particularly interested in the impact of synaptic input currents in CA1 *stratum radiatum*, we introduced electrophysiological parameters as defined in the NEURON CA1 neuron model of Combe et al.^17^. We generated an individual neuron dynamics model for each reconstructed dendritic tree accounting for a leak current, two fast Hodgkin-Huxley-type Na+ currents (somato-dendritic and axonal), a delayed rectifier, two A-type K+ currents (proximal and distal), an M-type K+ current, a mixed conductance hyperpolarization-activated h-current, a T-type Ca2+ current, an R-type Ca2+ current, two L-type Ca2+ currents (somatic and dendritic), a slow Ca2+-dependent K+ (SK) current and a Ca2+- and voltage-dependent K+ (BK) current.

To introduce synaptic currents, we accounted for the differences in spine density and length of the dendritic compartments between genotypes and conditions in order to position synaptic currents accordingly. Given that spine density was measured in 20 μm length dendritic segments, to obtain more realistic electrophysiological dynamics throughout dendritic branches, we accounted for the previously described linear decrease of synaptic weight from the trunk to the tips CA1 apical dendrites^61^. We scaled the weight of each synapse by the branch level order (summed path distance of all child branches to the root of the tree, https://www.treestoolbox.org/manual/LO_tree.html), which is minimal on the tips of the branches and maximal close to the trunk. Synaptic weight values were scaled by the level order in each point of the tree and normalized to maximal and minimal values of 0.6 and 0.4 spines per μm respectively based on Katz et al.^61^.

##### Averaged activity upon inhibitory feedback

To assess the impact of inhibition on the spiking frequency of CA1 neurons we extended our multi-compartmental models to include somatic inhibitory synapses. A pair of inhibitory neurons was simulated for each CA1 pyramidal cell to account for both inhibition of inhibitory and excitatory neurons^62^. Artificial integrate-and-fire inhibitory neurons were defined using the T2N IntFire2 structure. The number of inhibitory synapses was set based on known proportions of synapses in CA1 pyramidal and Parvalbumin neurons^63^. To explore how the averaged activity of different cells could coordinate their activity through inhibition, individual pyramidal cells were stimulated with regular 100Hz spike trains in 7.5% available synapses for 1 second. One multi-compartmental model was generated for each neuron. The somatic activity was measured in the model and spike times were recorded. The averaged activity power spectrum for each group was quantified by pooling the spike times of all the simulated neurons in a single array and obtaining its Fourier transform. The code for the multi-compartmental model simulations can be found at https://github.com/lmanubens/EE_CA1/tree/main/model.

#### Electrophysiological recordings

Field excitatory postsynaptic potentials (fEPSPs) were recorded in the *stratum radiatum* of the dorsal hippocampal CA1 region^52^ (Bregma, antero-posterior −1.06 to −2.54 mm; medio-lateral 0.5 to 1.6 mm), in response to stimulation of the Schaffer collateral pathway in WT and TG mice reared in NE conditions and after one month of EE conditions (number of slices/animals: WTNE = 11/5; TGNE = 11/5; WTEE = 9/5; TGEE= 16/5). The same analysis was performed in WT and TG mice reared under NE conditions for 4 months after termination of EE treatment to evaluate the long-term stability of EE effects (number of slices/animals: WTNE = 16/5; TGNE = 13/6; WTEEdis = 15/7; TGEEdis = 16/5). Following decapitation, the brain was quickly removed, placed on ice-cold cutting solution (in mM: KCl 2.5; MgSO4 3; NaHPO4 1.25; CaCl2 1; NaHCO3 26; sucrose 10) and gassed with 95% O2-5% CO2 to a final pH of 7.4. Four-hundred µm thick coronal slices were obtained with a vibratome (Leica). The slices were then placed in an interface style recording chamber (Fine Science Tools) and bathed in artificial cerebrospinal fluid (ACSF, in mM: NaCl, 124; KCl, 2.5; MgSO4, 1; NaHPO4, 1.25; CaCl2, 2.5; NaHCO3, 26; dextrose, 10), and then aerated with 95% O2-5% CO2 to a final pH of 7.4. Bath temperature was maintained at 32–34°C. Unfiltered recordings were obtained by means of glass electrodes (impedance 1-2 MΩ) through a Neurolog system (Digitimer) amplifier. Electrical stimuli were delivered with a concentric bipolar electrode (platinum-iridium) with the stimulus strength adjusted to a stimulation intensity that yielded a half-maximal response. For each slice, after determining a stable baseline, paired-pulse facilitation (PPF) was induced by a double-pulse (50 ms apart) stimulation protocol and represented as the ratio of both stimuli (fEPSP2/fEPSP1). After 15 min of baseline registering (pulse at 0.016 Hz), long-term potentiation (LTP) was induced via theta burst stimulation (TBS) (5 episodes at 0.1Hz; one episode was 10 stimulus trains of 4 pulses at 100 Hz; delivered at 5 Hz) and registered for 60 min (pulse at 0.016 Hz). Data are represented as the percentage of the baseline response. Recordings were digitized, acquired, and analyzed using a data acquisition interface, and software from Cambridge Electronic Design (Spike2; version 6.18).

#### Dyrk1A kinase activity assay

Dyrk1A kinase activity was measured in the hippocampus (number of animals: WTNE = 11; TGNE = 9; WTEE = 10; TGEE = 10 and after EE discontinuation: WTNE = 12; TGNE = 12; WTEE = 12; TGEE = 11) as previously described^10^. Briefly, samples were extracted in a Hepes lysis buffer and immunoprecipitated with a mouse anti-Dyrk1A antibody (Abnova RRID:AB_1574529; 3 μg/sample) immobilized on glutathione-sepharose beads (GE Healthcare). The recovered samples were analyzed by immunoblotting and *in vitro* kinase assays.

To determine the catalytic activity of Dyrk1A, 500 μg of the purified protein were incubated for 50 min at 30°C in 30 μl of phosphorylation buffer containing 2mM of specific Dyrk1A peptide^64^ [DYRKtide-RRRFRPASPLRGPPK], 1mM ATP, and [γ-32P]ATP (2 μCi/sample). Five μl of reaction aliquots were dotted onto P81 Whatman paper. After washing extensively with 5% phosphoric acid, counts were determined in a liquid scintillation counter. Relative kinase activity was obtained, normalizing against the amount of Dyrk1A protein present in the immunocomplexes. Immunocomplexes were resolved by 7.5% SDS-PAGE, transferred onto a nitrocellulose membrane and incubated with anti-Dyrk1A antibody (1/1000) overnight at 4°C. After that, membranes were incubated for 1 hour at RT with the corresponding secondary antibody. Detection was performed using ECL (GE Healthcare) and determined with LAS-3000 image analyzer (Fujifilm Image Reader V2.2; Fuji PhotoFilm). Protein quantification was performed using Image Gauge software version 4 (Fuji PhotoFilm).

#### Quantification and statistical analysis

Data were plotted as mean ± S.E.M. or ± S.D. and analyzed using the multivariate analysis of variance (MANOVA) for the statistical analysis of four groups containing two independent variables (genotype and treatment). Bonferroni-Holm was used for post-hoc analysis when a significant, or a trend to (p<0.09) genotype x treatment interaction was found. For those experiments with repeated events, comparisons were performed using MANOVA repeated measures. For structural analysis data we used an unbalanced block design, which allows an unequal number of observations per group (in our case, numbers of neurons reconstructed per genotype and/or treatment). In this case we used a mixed-effect linear model with the residual maximum likelihood (REML) method and blocked for biological replicates. P-values were obtained from the adjusted denominator degrees of freedom for linear estimates and t distributions calculated using the Kenward-Roger approximation (mixed function in the afex R package, version 0.28-1), adjusting pairwise comparisons with Bonferroni-Holm as pot-hoc. All the analyses were performed using R (version 4.1.0) and its packages stats (version 4.1.0) and emmeans (version 1.6.1). Results were considered significant when p<0.05 and trends were considered when p<0.1. All the metrics and analyses are available in an in-house developed Shiny App (version 1.6.0; https://linusmg.shinyapps.io/EE_TgDyrk1A/).

---

**Supplementary Figure 1.**
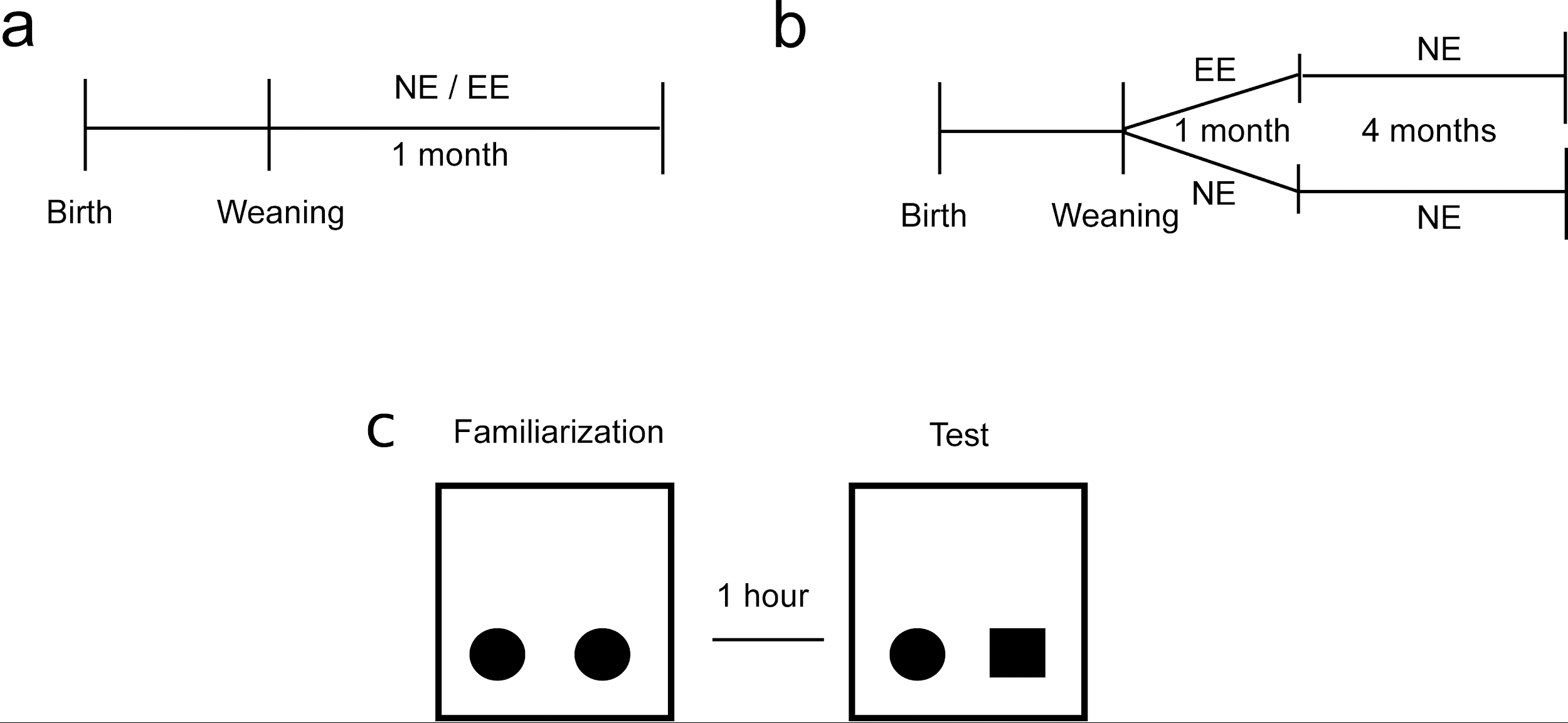
Experimental schedule used in wild type (WT) and TgDyrk1A (TG) mice for non-enriched (NE) and enriched (EE) rearing conditions. **b** Experimental schedule used to assess long-term effects of EE in WT and TG mice. Briefly, mice were reared for one month under NE or EE conditions followed by four months under NE conditions. **c** Schematic representation of the novel object recognition (NOR) paradigm.

**Supplementary Video 1.** Animation showing the dorsal CA1 region analyzed in this study.

**Supplementary Figure 2.**
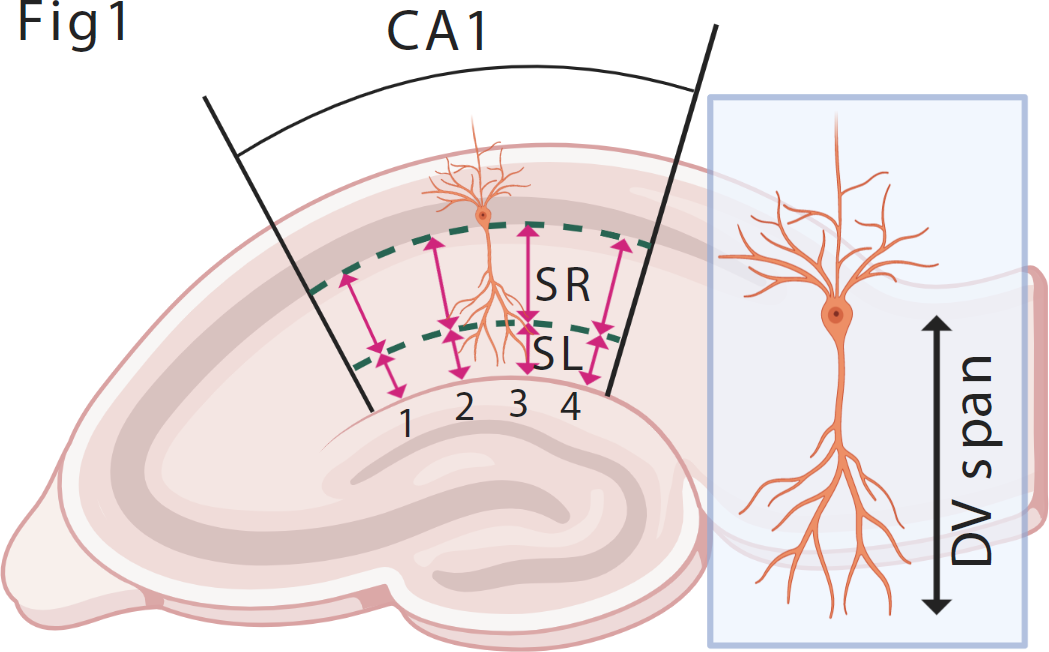
Neuromorphological analysis of dorsal hippocampal CA1 layer. The width of the stratum radiatum (*stratum radiatum*; intercept between the stratum pyramidale-radiatum border and the radiatum-lacunosum border) and stratum lacunosum (*stratum lacunosum*; intercept between the stratum radiatum-lacunosum border and the hippocampus sulcus), where apical dendrites of CA1 pyramidal neurons are located, were analyzed at four medio-lateral locations^52^ (Bregma, antero-posterior = −1.06 to −2.54 mm; medio-lateral 0.5 to 1.6 mm). The four locations were equispaced by ∼200µm from proximal to distal along the medio-lateral axis.

**Supplementary Figure 3.**
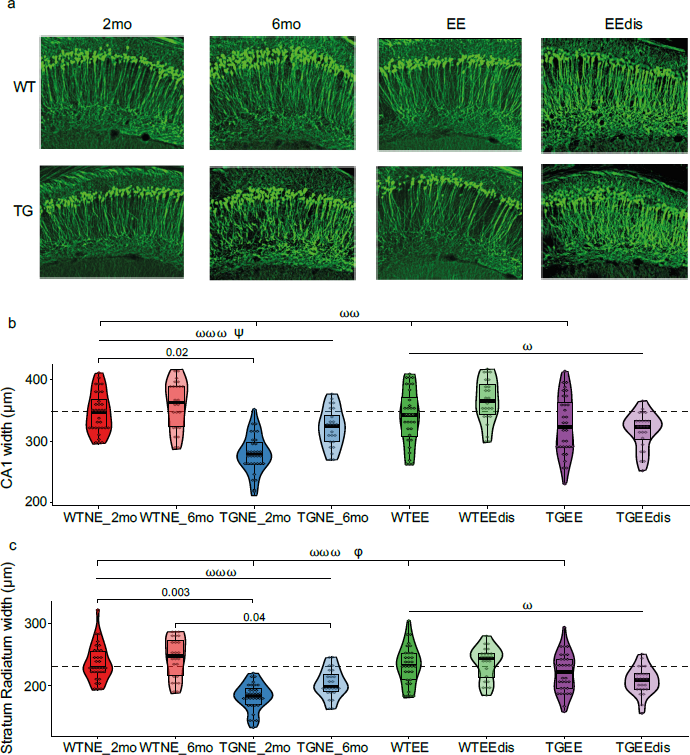
**a** Representative confocal micrographs showing YFP+ CA1 pyramidal neurons in WT and TG mice reared under NE at 2 (1^st^ column) and 6 (2^nd^ column) months of age, or one month EE (3^rd^ column) conditions and four months after EE discontinuation (EEdis; 4^th^ column). **b** and **c** Width of whole CA1 (b) and CA1 *stratum radiatum* layer (c) in WTNE and TGNE, at 2 and 6 months of age, WTEE and TGEE (2 months of age) and WTEEdis and TGEEdis four month after EE discontinuation (6 months of age). Data are presented as an overlay of dotplots, boxplots and violin plots. The dotted line indicates the median value of the WTNE (2mo) group. Number of slices/animals: WTNE (2mo) = 14/5; WTNE (6mo) = 24/8; TGNE (2mo) = 8/4; TGNE (6mo) = 9/4; WTEE (2mo) = 17/6; WTEEdis (6mo) = 18/6; TGEE (2mo) = 8/3; TGEEdis (6mo) = 19/7. Linear mixed effects: genotype effect ⍵ p<0.05; ⍵⍵ p<0.01; ⍵⍵⍵ p<0.001; genotype x age interaction ψ p<0.05; genotype x treatment interaction φ p<0.05.

**Supplementary Figure 4.**
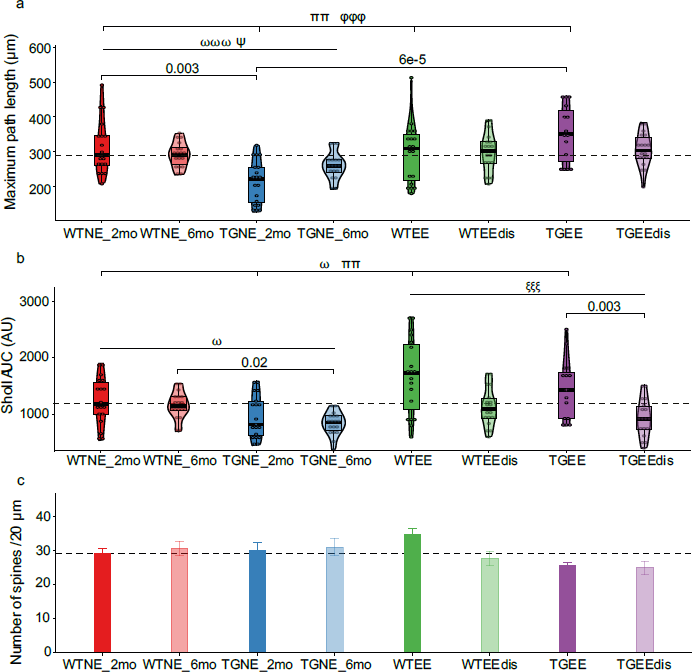
**a** Maximum path length of the dendritic trees. Data are presented as an overlay of dotplots, boxplots and violin plots. The dotted line indicates the median value of the WTNE (2mo) group. Values are given for WTNE and TGNE, at 2 and 6 months of age, WTEE and TGEE (2 months of age) and WTEEdis and TGEEdis four month after EE discontinuation (6 months of age). Number of neurons/animals WTNE (2mo) = 18/4; WTNE (6mo) = 18/5; TGNE (2mo) = 19/4; TGNE (6mo) = 18/5; WTEE (2mo) = 19/4; WTEEdis (6mo) = 18/3; TGEE (2mo) = 15/3; TGEEdis (6mo) = 18/4. **b** Area under the curve of the Sholl analysis distributions. Data are presented as an overlay of dotplots, boxplots and violin plots. The dotted line indicates the median value of the WTNE (2mo) group. Values represent AUC in WTNE and TGNE, at 2 and 6 months of age, WTEE and TGEE (2 months of age) and WTEEdis and TGEEdis four month after EE discontinuation (6 months of age). Number of neurons/animals: WTNE (2mo) = 18/4; WTNE (6mo) = 18/5; TGNE (2mo) = 19/4; TGNE (6mo) = 18/5; WTEE (2mo) = 19/4; WTEEdis (6mo) = 18/3; TGEE (2mo) = 15/3; TGEEdis (6mo) = 18/4. **c** Spine density (number of spines in a 20 μm segment of the dendrite). Data are expressed as mean ± SEM. The dotted line indicates the mean value of the WTNE (2mo) group. Number of dendrite segments/neurons/animals analyzed: WTNE (2mo) = 19/19/4; WTNE (6mo) = 20/20/4; TGNE (2mo) =16/12/4; TGNE (6mo) = 19/19/5; WTEE (2mo) = 19/19/4; WTEEdis (6mo) = 18/18/3; TGEE (2mo) = 16/9/4; TGEEdis (6mo) = 18/18/6. Linear mixed effects: genotype effect ⍵ p<0.05; ⍵⍵⍵ p<0.001; treatment effect ππ p<0.01; genotype x age interaction ψ p<0.05; treatment discontinuation effect ξξξ p<0.001; genotype x treatment interaction φφφ p<0.001.

**Supplementary Figure 5.**
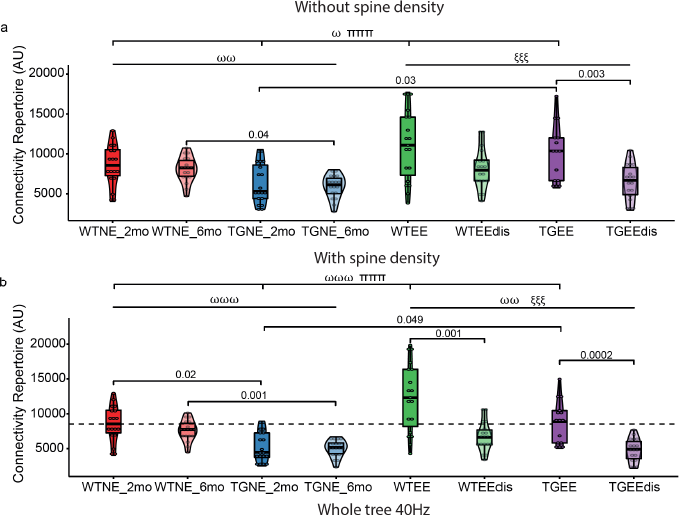
Connectivity repertoire of the CA1 apical trees of 2mo WT and TG reared under NE or EE conditions accounting for dendritic tree complexity and span differences (**a**) or also including spine density measurements (**b**). Data is presented as an overlay of dotplots, boxplots and violin plots. The dotted line indicates the median value of WTNE. Neurons/animals WTNE (2mo) = 18/4; WTNE (6mo) = 18/5; TGNE (2mo) = 19/4; TGNE (6mo) = 18/5; WTEE (2mo) = 19/4; WTEEdis (6mo) = 18/3; TGEE (2mo) = 15/3; TGEEdis (6mo) = 18/4. Linear mixed effects genotype effect ⍵ p<0.05, ⍵⍵ p<0.01, ⍵⍵⍵ p<0.001; treatment effect πππ p<0.001; treatment discontinuation effect ξξ p<0.01, ξξξ p<0.001.

**Supplementary Figure 6.**
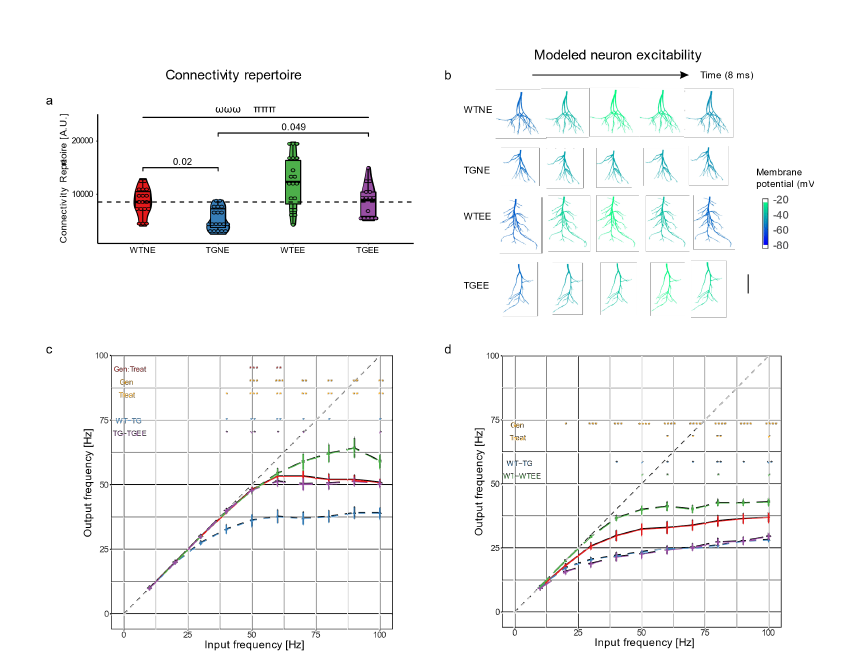
Connectivity repertoire and input-output frequency deficits in TgDyrk1A mice are partially rescued by environmental enrichment. **a** Connectivity repertoire of CA1 apical trees of 2mo WT and TG reared under NE or EE conditions is presented as an overlay of dotplots, boxplots and violin plots. The dotted line indicates the median value of WTNE (2mo). Neurons/animals: WTNE (2mo) = 18/4; TGNE (2mo) = 19/4; WTEE (2mo) = 19/4; TGEE (2mo) = 15/3. Linear mixed effects genotype effect ⍵⍵⍵ p<0.001; treatment effect πππ p<0.001. b Color-coded representation of the membrane potential (mV) modeled in the multicompartmental simulations on a 2D projection of representative apical dendritic trees (*SR* and *SL*) of each experimental group every 2 msec (ms) (30-38 ms of traces in Fig. S7a). Note that TGNE (2mo) neurons do not reach depolarization, a deficit recovered by EE. Scale bar = 100 μm. c-d Mean IO relation of reconstructed apical trees at frequencies from 10 to 100 Hz upon stimulation of *SR and SL* (c) and only of *SR* (d). Data are presented as the average of all the simulated neurons for each experimental group (mean ± SEM). Linear mixed effects, * p<0.05, ** p<0.01, *** p<0.001.

**Supplementary Figure 7.**
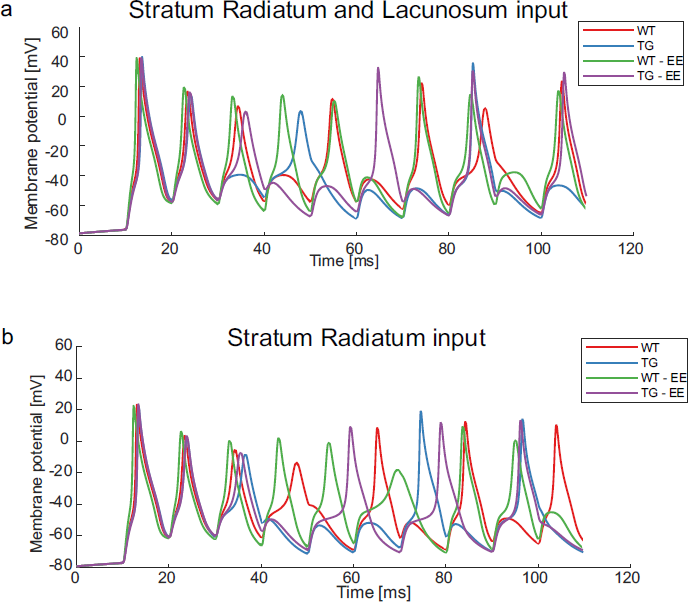
Somatic membrane potential of representative neurons obtained upon simulation of 100Hz input frequency on *stratum radiatum and lacunosum* (**a**) and on *stratum radiatum* only (**b**).

**Supplementary Figure 8.**
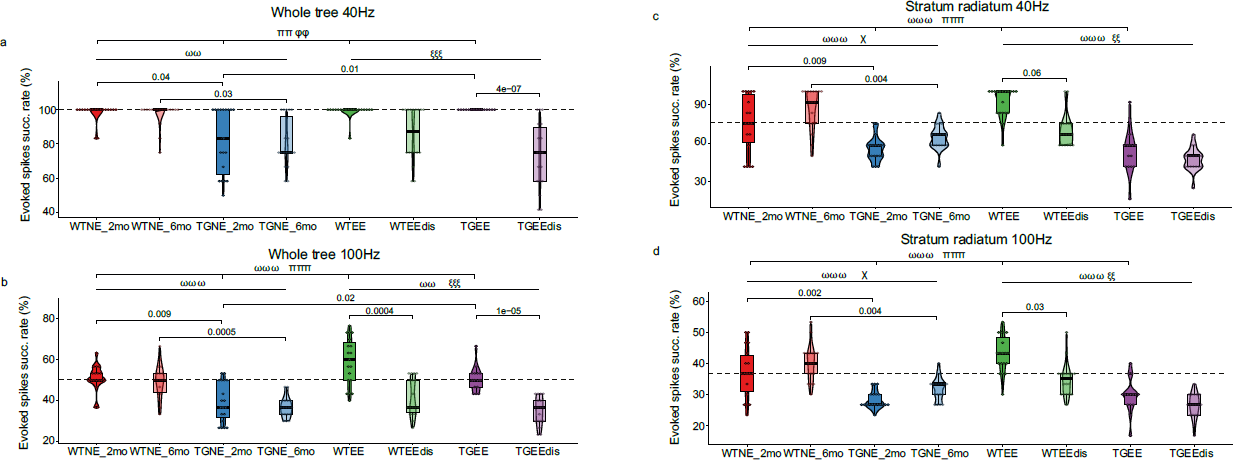
Success rate of action potentials evoked upon low-frequency stimulation on *stratum radiatum and lacunosum* (**a**) and only on *stratum radiatum* (**c**). Success rate of evoked action potentials upon high-frequency on *stratum radiatum and lacunosum* (**b**) and only on *stratum radiatum* (**d**). Data are presented as an overlay of dotplots, boxplots and violin plots. Dotted line indicates the median value of the 2mo WTNE group. Neurons/animals WTNE (2mo) = 18/4; WTNE (6mo) = 18/5; TGNE (2mo) = 19/4; TGNE (6mo) = 18/5; WTEE (2mo) = 19/4; WTEEdis (6mo) = 18/3; TGEE (2mo) = 15/3; TGEEdis (6mo) = 18/4. Linear mixed effects genotype effect ⍵⍵ p<0.01, ⍵⍵⍵ p<0.001; treatment effect π p<0.05, ππ p<0.01, πππ p<0.001; treatment discontinuation effect ξξ p<0.01, ξξξ p<0.001; genotype x treatment interaction φφ p<0.01; age effect χ p<0.05.

**Supplementary Figure 9.**
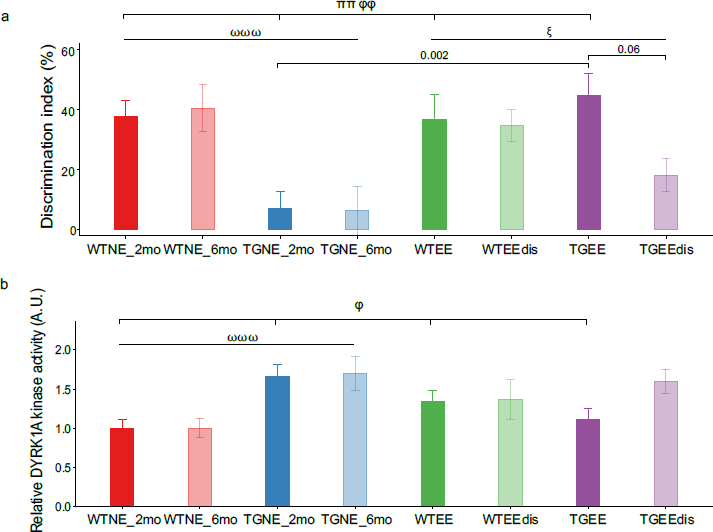
Long term effects of EE on cogntive performance and DRYK1A kinase activity are lost upon discontinuation. **a** Percentage of discrimination index during the test session of the NOR paradigm in WT and TG mice reared on EE and reared on EE followed by four months under NE conditions. Number of animals: WTNE-2mo = 14; TGNE-2mo= 15; WTEE (2mo) = 16; TGEE (2mo) = 19; WTNE-6mo = 11; TGNE-6mo = 12; WTEEdis (6mo) = 18; TGEEdis (6mo) = 18). **b** Relative DYRK1A kinase activity in the hippocampus. Number of animals: WTNE-2mo = 11; TGNE-2mo= 9; WTEE = 10; TGEE = 10; WTNE-6mo = 12; TGNE-6mo = 12; WTEEdis = 12; TGEEdis = 11. Data are expressed as mean ± SEM. Three-way ANOVA, Bonferroni-Holm as post-hoc indicated with numbers. genotype effect ⍵⍵⍵ p<0.001; treatment effect ππ p<0.01; genotype x treatment interaction φ p<0.05, φφ p<0.01.

**Supplementary Figure 10.**
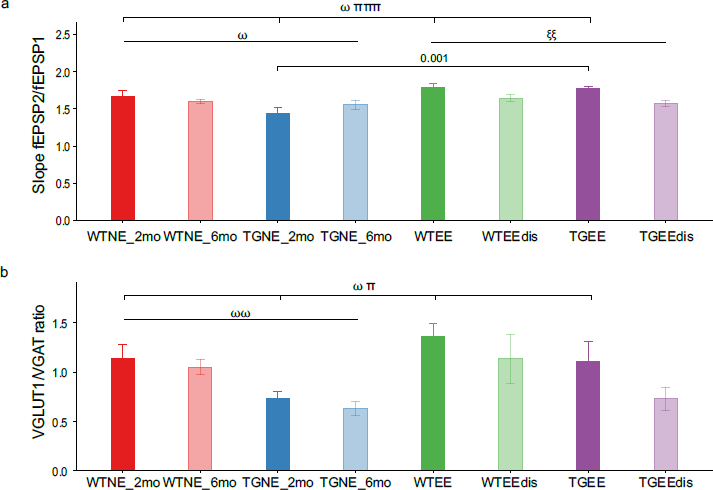
**a** Paired pulse facilitation (fEPSP2/fEPSP1) in the hippocampus with an inter-stimulus interval of 50ms (slices/animals: WTNE (2mo) = 11/5; TGNE (2mo) = 11/5; WTEE (2mo) = 9/5; TGEE (2mo) = 16/5; WTNE (6mo) = 18/5; TGNE (6mo) = 14/6; WTEEdis (6mo) = 20/7; TGEEdis (6mo) = 12/5). Three-way ANOVA genotype effect ⍵ p<0.05; treatment effect πππ p<0.001; treatment discontinuation effect ξξ p<0.01. Significance in numbers indicate pairwise Bonferroni-Holm as post-hoc. **b** Histogram showing the ratio between VGLUT1+ and VGAT+ punctae. WTNE (2mo) n = 7; TGNE (2mo) n = 7; WTEE (2mo) n = 8; TGEE (2mo) n = 7; WTNE (6mo) n = 5; TGNE (6mo) n = 6; WTEEdis (6mo) n = 5; TGEEdis (6mo) n = 4. Data are represented as mean + SEM. Three-way ANOVA, genotype effect ⍵ p<0.05, ⍵⍵ p<0.01; treatment effect π p<0.05.

**Supplementary Figure 11.**
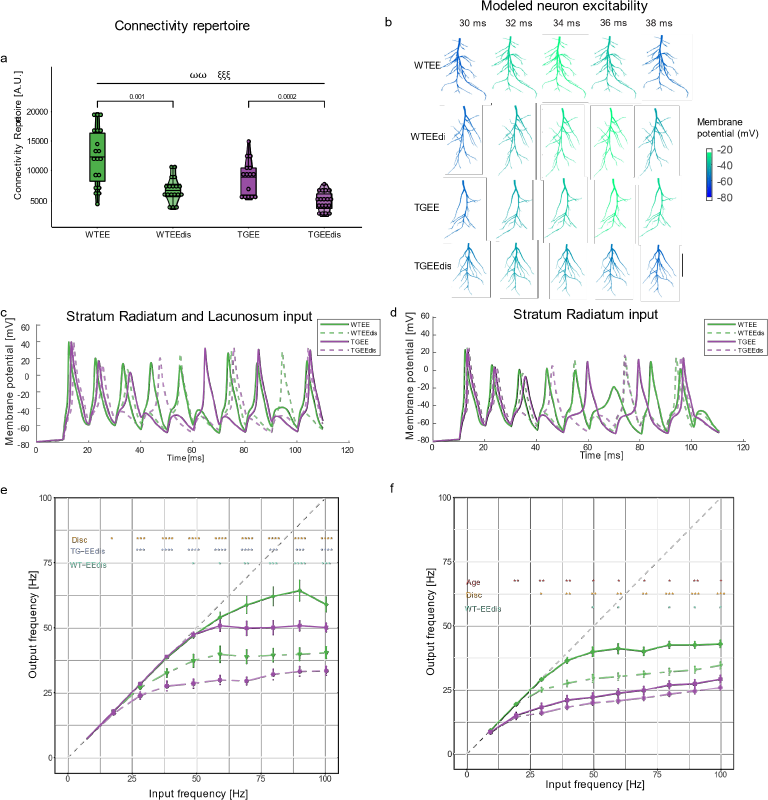
Connectivity repertoire and input-output frequency relationship after EE discontinuation. **a** Connectivity repertoire of CA1 apical trees obtained by Eq. 2. Data is presented as an overlay of dotplots, boxplots and violin plots. Number of neurons/animals analyzed: WTEE = 19/4; TGEE = 15/3; WTEEdis = 18/3; TGEEdis= 18/4. Linear mixed effects age x treatment interaction *** p<0.001. **b** 2D projection of representative apical dendritic trees of each experimental group seen in the coronal plane with a color-coded representation of the membrane potential modeled in the multicompartmental simulations of simultaneous synaptic stimulation of 15% of available synapses at 100 Hz. Scale bar = 100 μm. **c-d** Somatic membrane potential obtained in the simulation for simultaneous inputs in *SR* and *SL* (c) and *SR* (**d**) activated with 100Hz input in representative neurons. **e-f** IO frequency relation obtained in the multicompartmental model by introducing synaptic currents in 15% of available synapses in the reconstructed apical trees at frequencies from 10 to 100 Hz. Data is presented as the average of all the simulated neurons for each experimental group (mean ± SEM). Linear mixed effects, * p<0.05, ** p<0.01, *** p<0.001.

**Supplementary Table 1.**
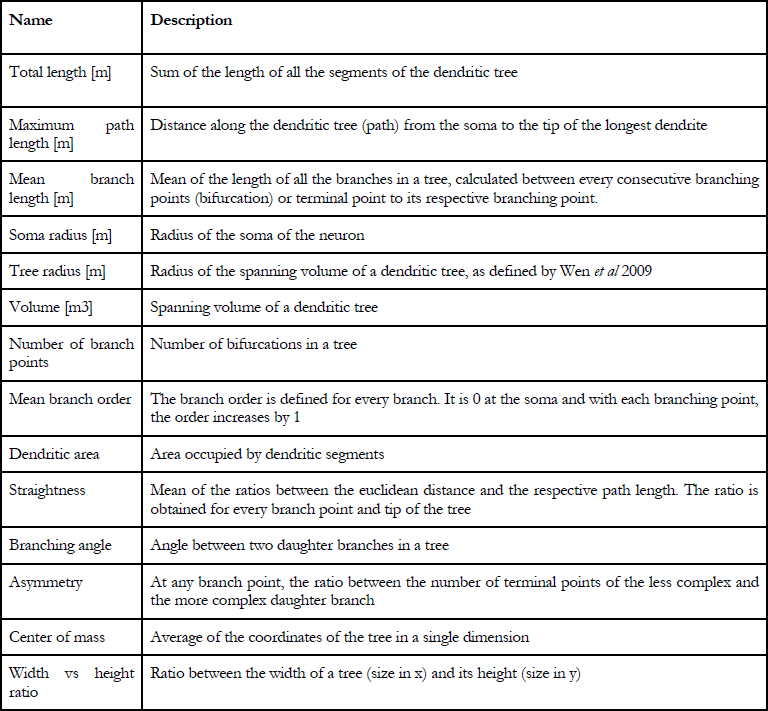
Description of the selected neuromorphological metrics. Each metric has one value for the whole tree.

| Name | Description |
| --- | --- |
| Total length [m] | Sum of the length of all the segments of the dendritic tree |
| Maximum path length [m] | Distance along the dendritic tree (path) from the soma to the tip of the longest dendrite |
| Mean branch length [m] | Mean of the length of all the branches in a tree, calculated between every consecutive branching points (bifurcation) or terminal point to its respective branching point. |
| Soma radius [m] | Radius of the soma of the neuron |
| Tree radius [m] | Radius of the spanning volume of a dendritic tree, as defined by Wen *et al* 2009 |
| Volume [m<sup>3</sup>] | Spanning volume of a dendritic tree |
| Number of branch points | Number of bifurcations in a tree |
| Mean branch order | The branch order is defined for every branch. It is 0 at the soma and with each branching point, the order increases by 1 |
| Dendritic area | Area occupied by dendritic segments |
| Straightness | Mean of the ratios between the euclidean distance and the respective path length. The ratio is obtained for every branch point and tip of the tree |
| Branching angle | Angle between two daughter branches in a tree |
| Asymmetry | At any branch point, the ratio between the number of terminal points of the less complex and the more complex daughter branch |
| Center of mass | Average of the coordinates of the tree in a single dimension |
| Width vs height ratio | Ratio between the width of a tree (size in x) and its height (size in y) |

## SUPPLEMENTARY TEXT

### Input-output relationship when stimulating only *Stratum Radiatum*

With stimulation of only *SR*, WT neurons responded to both LF and HF inputs with an output firing rate corresponding to low γ (30-40 Hz). The ability of CA1 to switch to low γ output when it receives inputs from only CA3 is consistent with phase-locking with CA3 and low γ synchronization during exploration of a familiar environment^34^. In 2mo TGNE neurons the firing rate was in the β range (22-28Hz) (Fig. S6d; IO frequency TGNE (2mo) vs WTNE (2mo) p<0.01 for HF inputs). For a detailed plot of success rates of evoked spikes see Fig. S8c (LF; linear mixed effects genotype effect for 2mo groups F(1,35.31)= 46.52, p=6e-8; TGNE (2mo) vs WTNE (2mo) Bonferroni-Holm as post-hoc: p=0.009) and S8d (HF; linear mixed effects genotype effect for 2mo groups F(1,31.38)= 52.42, p=4e-8; TGNE (2mo) vs WTNE (2mo) Bonferroni-Holm as post-hoc: p=0.002). Our results suggest that 2mo TGNE neurons are not capable of evoking γ upon *SR* stimulation. When modelling stimulation only to SR, WTEE neurons slightly increased their ability to respond to both LF and HF inputs with an output firing rate still corresponding to low-γ (30-40 Hz), while upon LF stimulation TGEE neurons kept their output at the β band (<30Hz) (Fig. S6d linear mixed effects treatment effect p<0.05; WTEE vs WTNE p<0.05; Fig. S8c success rate of evoked spikes upon LF stimulation of only SR linear mixed effects treatment effect for 2mo groups N.S.; Fig. S8d HF stimulation of only SR linear mixed effects treatment effect for 2mo groups N.S.).

After EE discontinuation, upon selective simulation on only SR, both WT and TG neurons had a slight reduction in the output firing rate. Thus, both LF and HF inputs, showed reduced firing rates in WTEEdis (6mo) neurons, reaching lower firing rate than WTEE (2mo) only at the lower limit of low-γ at 100Hz stimulation. TGEEdis (6mo) showed a non-significant reduction in the firing rate still at the β band in comparison to TGEE (2mo) (Fig. S11f; IO frequency discontinuation effect p<0.01; Fig. S8a linear mixed effects discontinuation effect F(1,60.18)= 10.31 p=0.002; WTEEdis (6mo) vs. WTEE (2mo), p=0.06; Figs. 7d and S8b; linear mixed effects discontinuation effect F(1,53.19)= 12.89 p=7e-4; WTEEdis (6mo) vs. WTEE (2mo), p=0.03). When we tested for the age effect on NE groups we found increased success rate of evoked spikes with LF and HF inputs at 6mo (Fig. S8a, S8b and S8f; age effect with LF F(1,93.77)=8.61, p=4e-3 and HF input F(1,82.28)=5.93, p=0.02, IO frequency age effect p<0.05), which was not significant for genotype pairwise comparisons. Regardless the age-related increase in success rate of evoked spikes, no significant differences were found in EEdis vs 6mo NE groups, suggesting that the effects of a short treatment with EE on SR input integration have no impact in the long term (Fig. S8a and S8b).

