## Supplementary material for "Deficits in neuronal architecture but not over-inhibition are main determinants of reduced neuronal network activity in a mouse model of overexpression of *Dyrk1A*": Resource Table

### KEY RESOURCES TABLE

| REAGENT or RESOURCE | SOURCE | IDENTIFIER |
| --- | --- | --- |
| Antibodies | | |
| Polyclonal antibody rabbit IgG fraction anti-Lucifer yellow | Life Technologies - Thermo Fisher Scientific | RRID:AB_2536190 |
| Secondary antibody goat anti-rabbit IgG 488 | Invitrogen - Thermo Fisher Scientific | RRID:AB_2576217 |
| Mouse anti-Dyrk1A antibody | Abnova | RRID:AB_1574529 |
| Chemicals, Peptides, and Recombinant Proteins | | |
| Dyrk1A peptide | Sigma | SRP0678 |
| Lucifer Yellow | Sigma | L 0259 |
| Deposited Data | | |
| Neuron reconstructions | NeuroMorpho.org | N/A |
| Experimental Models: Organisms/Strains | | |
| Mouse:Thy1:B6.Cg-Tg(Thy1-YFPH)2Jrs/J | The Jackson Laboratories | 003782 |
| Mouse:WT:C57BL/6JXSJL | Charles River | N/A |
| Mouse:WT:C57BL/6JXSJL-TgDyrk1A | Dierssen lab, 4 | N/A |
| Software and Algorithms | | |
| SMART | Panlab | N/A |
| Fiji | 13 | N/A |
| NeuTube | 16 | N/A |
| Trees Toolbox | 17 | N/A |
| Imaris | Bitplane | N/A |
| Huygens | Scientific Volume Imaging | N/A |
| T2N | 21 | N/A |
| Connectivity repertoire estimation | this paper | Zenodo |
| CA1 pyramidal neuron model | 19 | NRNID 244416 |
| R | R Foundation for Statistical Computing | N/A |
| Spike2 | Cambridge Electronic Design | N/A |
| Shiny app for analysis | this paper | 10.5281/zenodo.7318051 |
